# Extraocular, rod-like photoreceptors in a flatworm express xenopsin photopigment

**DOI:** 10.1101/538538

**Authors:** Kate A. Rawlinson, François Lapraz, Johannes Girstmair, Fraser Simpson, Katharine E. Criswell, Mark Terasaki, Edward R. Ballister, Miklos Boldogkoi, Richard J. McDowell, Jessica Rodgers, Brian K Hall, Robert J. Lucas, Maximilian J. Telford

## Abstract

Animals detect light using opsin photopigments. One recently classified opsin clade, the xenopsins, found in lophotrochozoans, challenges our views on opsin and photoreceptor evolution. Originally thought to belong to the G*α*_i_-coupled ciliary opsins, xenopsins are now understood to have diverged from ciliary opsins in pre-bilaterian times, but little is known about the cells that deploy these proteins, or if they form a photopigment and drive phototransduction. We characterized xenopsin in a flatworm, *Maritigrella crozieri,* and found that it is expressed in a larval eyespot, and in an abundant extraocular cell type around the adult brain. These distinct cells house hundreds of cilia in an intra-cellular vacuole (a phaosome). Cellular assays show *Mc* xenopsin forms a photopigment and couples to Gα _i/o_ in response to light. These findings reveal a novel photoreceptor cell type and opsin/G-protein couple, and highlight the convergent enclosure of photosensitive cilia in flatworm phaosomes and jawed vertebrate rods.

## Introduction

Light is a key biological stimulus for most animals, and a rich diversity of photosensitive cells has evolved. Depending on the form of their elaborated apical plasma membranes, these cells have been characterized as either ciliary photoreceptors (CPRs) or rhabdomeric (microvillar) photoreceptors (RPRs) (Eakin, 1982). When rhabdomeric and ciliary photoreceptors coexist in the same organism, one type (rhabdomeric in most invertebrates, ciliary in vertebrates) typically dominates in the eyes while the other performs nonvisual functions in the eyes or is present as extraocular photoreceptors (Arendt et al., 2004; Yau and Hardie, 2009). Photopigments are responsible for the light-dependent chemical reactions in these cells, and all animal phyla, with the exception of sponges, employ photopigments composed of opsin-class G-protein-coupled receptors (GPCRs) coupled with a light-sensitive chromophore (a retinaldehyde) (Nilsson, 2013; Bok et al., 2017). The initial characterization of opsins in bilaterian animals identified several conserved opsin gene families (Terakita, 2005), and each family has been associated with distinct photoreceptor cell types and specific downstream G-protein phototransduction cascades (reviewed in Lamb, 2013). For example, ciliary (c)-opsins are expressed in ciliary photoreceptor cells where they typically activate members of the Gα _i_ families, while rhabdomeric (r)-opsins activate Gα _q_ family members and are expressed in rhabdomeric photoreceptors (Shichida and Matsuyama, 2009). The recent accumulation of sequence data from a taxonomically broader set of animals has, however, revealed a far greater diversity of opsins (Porter et al., 2012; Ramirez et al., 2016; Bok et al., 2017; Vöcking et al., 2017), and the ability to localise the opsin mRNA transcripts and proteins in a diversity of animals has uncovered many new and morphologically divergent photosensitive cell types, both ocular and extraocular (Vöcking et al., 2017; Bok et al., 2017).

The recent identification of one novel group of opsins – the xenopsins (Ramirez et al., 2016) – is leading to the reevaluation of eye and photoreceptor cell type evolution (Vöcking et al., 2017). Xenopsins have been found in several lophotrochozoan phyla: molluscs, rotifers, brachiopods, flatworms and an annelid (Ramirez et al., 2016; Vöcking et al., 2017). They share with some ciliary opsins a characteristic c-terminal sequence motif (NVQ) and were originally classified as part of the c-opsins (Passamaneck et al., 2011; Albertin et al., 2015; Yoshida et al.. 2015). All recent opsin phylogenies have, however, shown xenopsins to be phylogenetically distinct from c-opsins (Ramirez et al., 2016; Bok et al., 2017; Vöcking et al., 2017; Quiroga Artigas et al., 2018) and gene structure analysis also supports this distinction (Vöcking et al., 2017). The relationship between xenopsins (lophotrochozoan protostome specific) and c-opsins (which are found in protostomes and deuterostomes) suggests that both opsins were present in the last common ancestor of Bilateria, and that xenopsins were subsequently lost in deuterostomes and ecdysozoan protostomes (Vöcking et al., 2017). Existing data on the expression of xenopsins are limited to the larval stages of a chiton and a brachiopod. In the brachiopod, xenopsin is expressed in cells with elaborated cilia and shading pigment i.e. pigmented eyespots (Passamaneck et al., 2011), whereas, unusually, in the chiton larva it is co-expressed with r-opsin in cells containing both cilia and microvilli. Some of these cells are supported by pigmented cells i.e. they probably form simple eyes, whereas others lack pigment and cannot act as visual photoreceptors (Vöcking et al., 2017).

While the presence of xenopsins in cells with elaborated ciliary surfaces and their association with pigmented cells is strongly suggestive of a role for xenopsins in photoreception, this function has not yet been demonstrated. Furthermore, if xenopsins do detect light, the subsequent phototransduction pathway is unknown. Determining these factors, and better understanding the phylogenetic distribution of xenopsins and of the cells in which they are expressed is essential for understanding the evolution of this opsin subtype and of the photoreceptors that use them (Arendt, 2017).

Flatworms (Platyhelminthes) are one of the most diverse and biomedically important groups of invertebrates. Their eyes typically consist of photoreceptors with rhabdomes of microvilli that are associated with pigmented shading cells (Sopott-Ehlers et al., 2001) and which express rhabdomeric opsin (Sanchez and Newmark, 1999) and conserved members of the r-opsin signaling cascade (e.g. Gα _q_, Trp channel-encoding genes) (Lapan and Reddien, 2012). The presence and nature of ciliary photoreceptors in flatworms is still unclear but the description of xenopsins (but not c-opsins) in flatworms (Vöcking et al., 2017) suggests CPRs may exist. Furthermore, ultrastructural studies have identified cells with elaborated ciliary membranes - putative CPRs (Sopott-Ehlers, 1991; Lyons, 1972, Kearn, 1993, Rohde and Watson, 1991) - but these have not been studied at the molecular level. In larvae of the polyclad *Pseudoceros canadensis*, ultrastructural studies identified three different types of CPR; the epidermal eyespot – a pigmented epidermal cell with elaborated ciliary membranes (Lanfranchi et al., 1981; Eakin and Brandenburg, 1981); a cerebral eye consisting of a CPR adjacent to two RPRs cupped by a supporting pigmented cell (Eakin and Brandenburg, 1981); and distinct extraocular cells in the epidermis possessing multiple cilia projecting into an intra-cellular vacuole (Lacalli, 1983) known as a phaosome (Fournier, 1984). This phaosomal cell type is found in all classes of flatworm (except triclads and bothrioplanids)(Sopott-Ehlers et al., 2001; Fournier, 1984), and the distinct morphology led to the suggestion that they are a derived feature of flatworms (Sopott-Ehlers et al., 2001).

Here we analyse xenopsin expression in a polyclad flatworm, in both larval and adult stages. We explore whether polyclad xenopsin can form a photopigment, and which class(es) of G-protein it can couple to by carrying out the first functional cellular assays on a xenopsin. For the first time we record the presence of xenopsin expressing cells in an adult lophotrochozoan, and characterize the ultrastructure of these unusual ciliary phaosome cells demonstrating similarities to jawed vertebrate rod photoreceptors. Together our findings show that xenopsin forms a photopigment driving phototransduction through G_i/o_ signalling, and it’s expression in ciliary phaosome cells establishes them as photoreceptors.

## Results

### 1. Xenopsins and rhabdomeric opsins in flatworms

A 346 amino acid gene product showing similarity to protostome c-opsin and xenopsin was predicted from a *Maritigrella crozieri* transcriptome contig using BLAST (Madden, 2002). Opsins showing similar degrees of similarity were found in transcriptomes from five other flatworm taxa (three polyclads; *Prostheceraeus vittatus, Stylochus ellipticus, Leptoplana tremellaris* and two triclad species *Schmidtea mediterranea, Dendrocoelum lacteum*). We did not find homologous sequences in the remaining 24 flatworm species representing other flatworm classes, including those in which putative CPRs have been described (catenulids, macrostomids, rhabdocoels, proseriates, monogeneans, cestodes and trematodes). Searching more broadly we found additional opsins similar to protostome c-opsins and xenopsins in a bryozoan, *Bugula nerita*, and in a chaetognath, *Pterosagitta draco.*

In our phylogenetic analyses of these putative flatworm, bryozoan and chaetognath opsins in the context of the metazoan opsin gene family, all cluster with xenopsins (**Figure 1**; **supplementary figure 1**). Several polyclad flatworm species show xenopsin paralogs distributed across two xenopsin subgroups (Vöcking et al., 2017); our six polyclad and triclad sequences all group with clade A and we add xenopsins from three additional taxa to this clade; *Maritigrella crozieri, Dendrocoelum lacteum, Leptoplana tremellaris* (**Figure 1**; **supplementary figure 1**). The xenopsins are a well-supported monophyletic group most closely related to cnidopsins. The xenopsin/cnidopsin group is sister to the tetraopsins and all are part of a larger clade including bathyopsins and canonical c-opsins (**Figure 1**).

**Figure 1.**
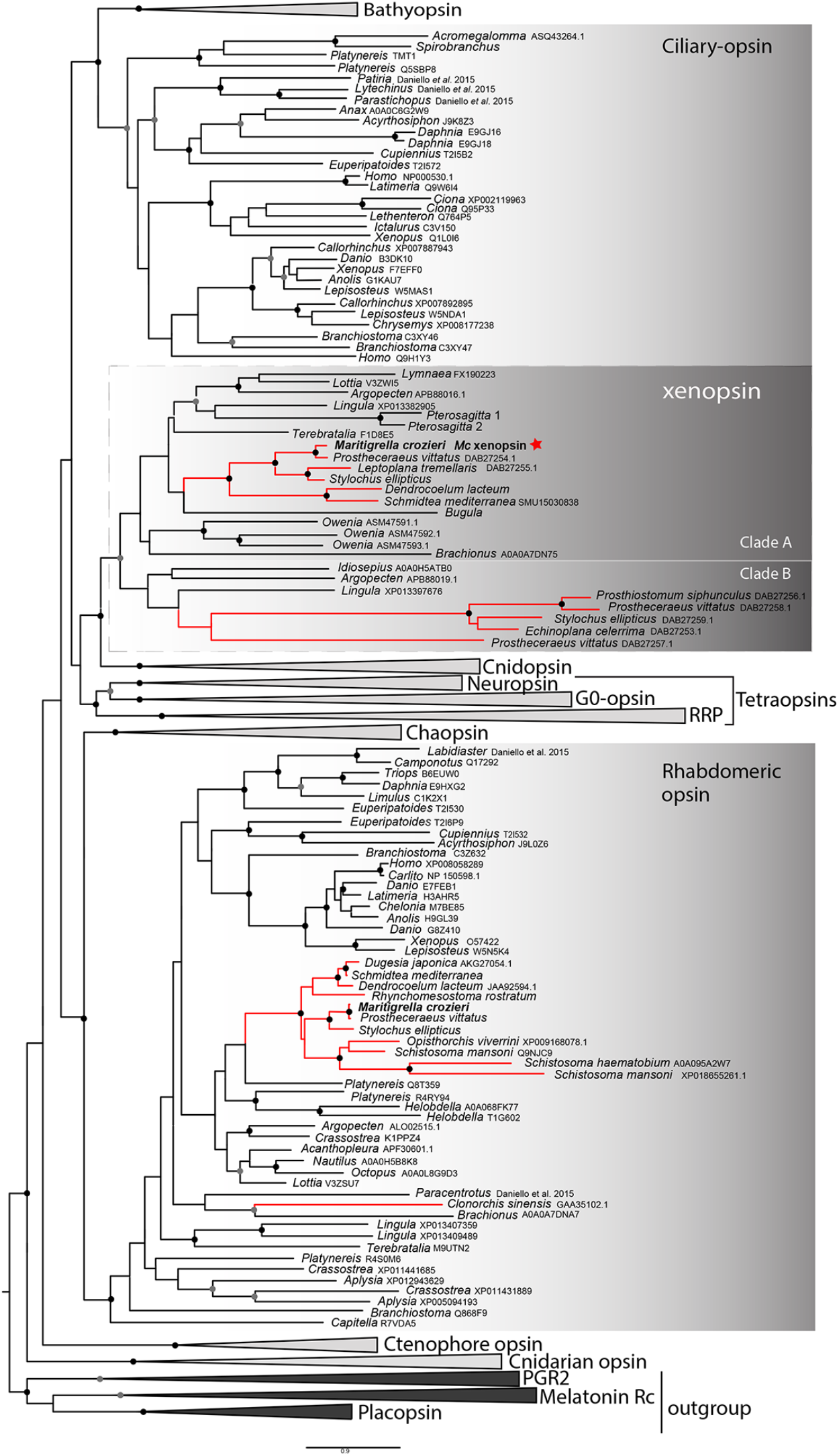
Phylogenetic analysis of metazoan opsins supports flatworm ciliary-like opsins as xenopsins, and confirms a clade of flatworm rhabdomeric opsins. Support for node is calculated using 1000 Ultrafast bootstrap replications as well as 1000 SH-aLRT replicates and approximate aBayes single Branch testing. Black dots indicate nodes with support values for 3 tests ≥ 95 (0.95 for SH-aLRT replicates). Grey dots indicate nodes with support values for 3 tests ≥ 90 (0.90 for SH-aLRT replicates). Scale bar unit for branch length is the number of substitutions per site. Branches in red correspond to flatworm opsin sequences. See **Supplementary figure 1** for uncollapsed tree and **Figure 1 – source data 1** for gene accession numbers. The new xenopsin sequences we found in polyclad and triclad flatworms, plus a bryozoan and chaetognath, all fall within clade A of the xenopsins.

Support for the relationships between these well-defined opsin subtypes is very low, indicating that these relationships should be interpreted cautiously. The need for caution is reinforced by the observation that removing the smaller opsin clades from our dataset (chaopsins, bathyopsins, ctenophore and anthozoan opsins), changes the topology of the deeper nodes of our trees (**Supplementary figure 2**).

The flatworm xenopsin protein sequences possess seven transmembrane domains (characteristic of all G protein-coupled receptors) as well as a conserved lysine in transmembrane domain VII, which is specific to opsins and which forms a Schiff base with the retinal chromophore to form a photopigment (**Supplementary figure 3**). *Mc* xenopsin also possesses a tripeptide motif, NxQ, at positions 310–312, which is reported in other xenopsins (Passamaneck et al., 2011, Vöcking et al., 2017) and in ciliary opsins where it is known to be crucial for G-protein activation (Marin, 2000, Gühmann et al., 2015).

We have found that a second amino acid signature, VxPx, found in vertebrate ciliary opsins at positions 423-426 is also present in xenopsins as well as in ciliary opsin sequences from non-vertebrate chordates (tunicate and lamprey), annelid c-opsins and cnidarian cnidopsins (**Supplementary figure 3**). In c-opsins this motif directly binds the small GTPase Arf4 to direct vertebrate rhodopsin (a ciliary opsin) to the primary cilia (Deretic et al., 2005). The presence of this motif in some ciliary opsins, xenopsins and cnidopsins suggests that Arf4 may be a shared mechanism for the active delivery of these opsins to the cilia in CPRs.

A 422 amino acid gene product related to rhabdomeric opsin was also predicted from a *Maritigrella crozieri* transcriptome contig. Nine more flatworm rhabdomeric-like opsins were predicted from both free-living and parasitic species. They possess a tripeptide motif (HP[K|R]) (supplementary figure 3) following the transmembrane helix VII, which is critical for G-protein binding in r-opsins (Plachetzki and Oakley, 2007). In our phylogenetic analysis, *Maritigrella r-opsin* and all flatworm *r-opsin* sequences (except one from the liver fluke *Clonorchis sinensis*) fall in a monophyletic group containing lophotrochozoan and ecdysozoan r-opsins, and deuterostome melanopsins (**Figure 1; Supplementary figure 1**).

Our reconstructions of the opsin gene family have resolved *Maritigrella* genes as orthologues of xenopsins and r-opsins (**Figure 1**), and we designated these genes as *Maritigrella crozieri xenopsin* (*Mc-xenopsin*) and *Maritigrella crozeri rhabdomeric opsin (Mc-r-opsin)*.

### 2. Mc xenopsin and r-opsin are expressed in distinct larval eyes

A pair of cerebral eyes containing rhabdomeric photoreceptors and three putative ciliary photoreceptor types (see Introduction) have been described in polyclad larvae (Lanfranchi et al., 1981; Eakin and Brandenburg, 1981; Lacalli, 1983). To identify these cells in *M. crozieri,* we used immunostaining with antibodies directed against *Mc* xenopsin and acetylated tubulin (which specifically labels stabilized microtubules in cilia and axons) to identify CPRs and *in situ* hybridization to determine the expression of *Mc r-opsin.*

*In situ* hybridization shows that both cerebral eyes house *r-opsin*^+^ photoreceptors (**Figure 2B & D**). Only one CPR type – the epidermal eyespot, shows xenopsin expression, however (**Figure 2B – Ci, supplementary videos 1 and 2**). In another polyclad this epidermal eyespot consists of two cells; a cup-shaped pigmented cell bearing flattened cilia and a distal cell that rests partly in the concavity of the pigmented cell (Eakin and Brandenburger, 1981). The epidermal eye develops before the cerebral eyes in *M.crozieri* (Rawlinson 2010) and xenopsin is expressed in this eyespot during embryogenesis (**Supplementary Figure 4**). The ontogenetic fate of this epidermal eyespot is unknown; it could be a transient larval character, as all pigmented eyes in adult *Maritigrella* are sub-epidermal and express *r-opsin* (Rawlinson *pers obs*) (**Figure 3I**). A cerebral eye with two rhabdomeric photoreceptors and one ciliary photoreceptor as reported from *Pseudoceros canadensis* (Eakin and Bradenburger, 1981), is not present in *Maritigrella* larvae as we show that neither of the two *r-opsin*^+^ rhabdomeric eyes expresses xenopsin. The third type of putative CPR are the unpigmented phaosomal CPRs described in the epidermis of a larval stage (Lacalli, 1983). Given the presence of acetylated tubulin^+^ cells in the dorsal epidermis of *Maritigrella* larvae, we think these are multi-ciliated phaosomal cells, but they did not express xenopsin (**Figure 2C & F**).

**Figure 2.**
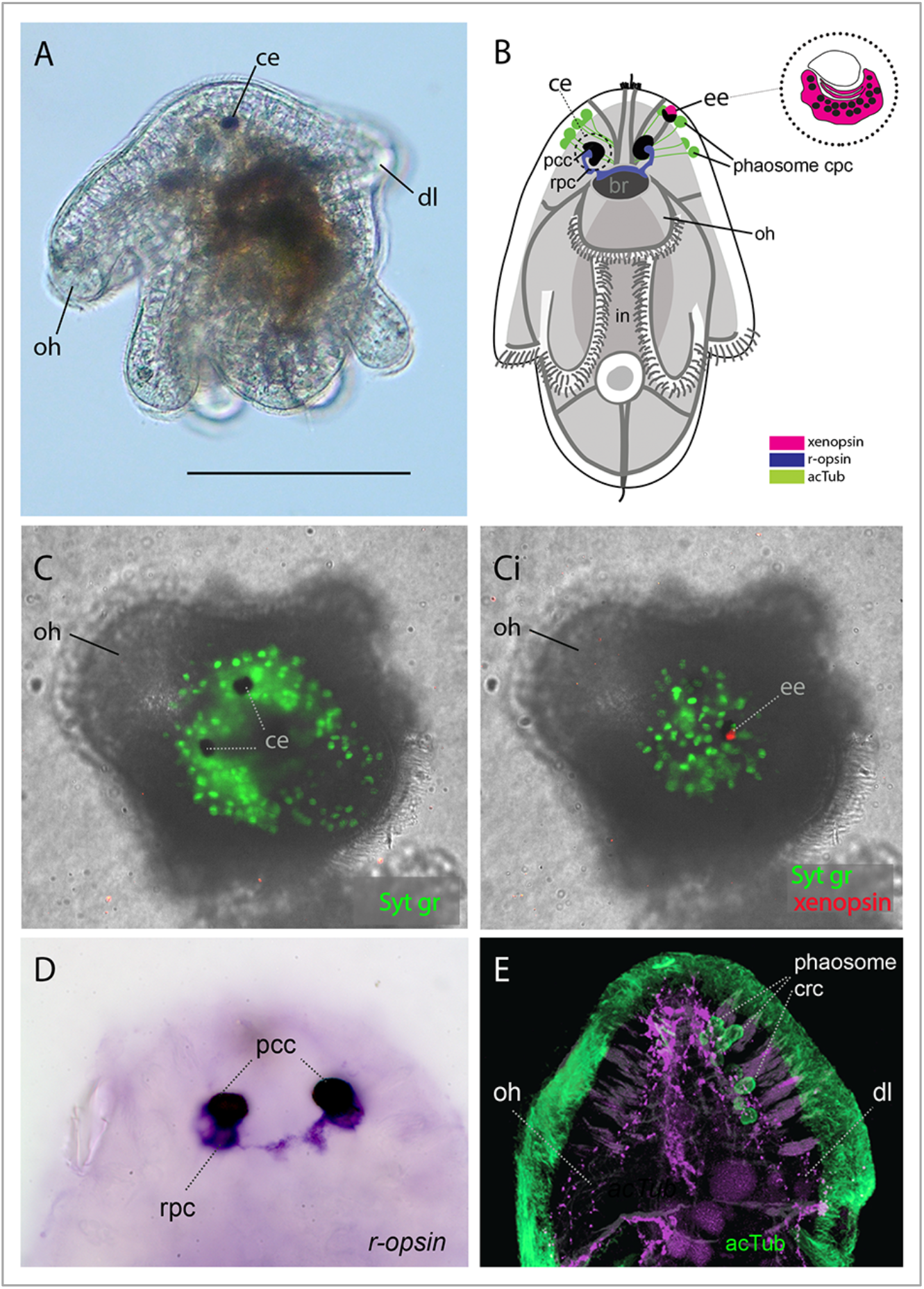
Xenopsin and rhabdomeric opsin expression in the larval eyes of *Maritigrella crozieri* (1-2 days posthatching). A) Left lateral view of live larva showing a pigmented cerebral eye (ce), the oral hood (oh) and dorsal lobe (dl)(scale = 150μm). B) A schematic of a larva showing the three putative ciliary photoreceptor cell (cpc) types: epidermal eye (ee), phaosome cpc, and cerebral eye (ce). C &Ci) Apical view of larvae showing xenopsin expression (red) in the epidermal eye (OpenSPIM images, Syt gr = Sytox green, staining nuclei and bright-field images also reveal photoreceptor pigments), D) both cerebral eyes (at 1 day post hatching) consist of an rhabdomeric opsin^+^ rhabdomeric photoreceptor cell (rpc) and a pigmented cup cell (pcc), E) the acTub^+^ phaosome cpc’s do not express xenopsin (see C). *in* = intestine, acTub = acetylated tubulin.

### 3. In adult Maritigrella, Mc xenopsin is expressed in extraocular ciliary phaosomal photoreceptors and Mc r-opsin is expressed in pigmented eyes

Immunofluorescence and *in situ* hybridization on sections of adult *Maritigrella* (**Figure 3**) show distinct xenopsin and *r-opsin* positive cells. To identify potential ciliary photoreceptors in adult *Maritigrella* we first used antibodies against acetylated tubulin and discovered two clusters of up to 100 acetylated tubulin^+^ cells, one either side of the brain (**Figure 3B, Bi & Di**). The cells are distributed from the anterior to the posterior of the brain (Fig.3Di) and extend laterally above nearby branches of the intestine (**Figure 3C**). Histological staining showed that these cells are embedded in extracellular matrix outside of, and lateral to, the brain capsule (**Figure 3C & Ci**) and that they sit in close proximity to the main nerve tracts (**Figure 3Ci**). The cells are stalked with a nucleus at one end and an intra-cellular vacuole (a phaosome) at the opposite end, into which multiple cilia protrude (Figure **3E & Ei**). *Mc* xenopsin was strongly co-expressed with acetylated tubulin in these cells (**Figure 3F-H**). These ciliated, phaosomal, xenopsin^+^ cells sit ventro-lateral to the *r-opsin*^+^ cells that extend from the pigmented cell cup down to the brain (**Figure 3I**). As is typical for r-opsin-expressing cells, they also express Gα _q_ as revealed by antibody staining (**Figure 3J**). The non-overlapping expression of xenopsin and *r-opsin* indicates that these opsins are expressed in two distinct photoreceptor types, with *r-opsin* expressed in the eyes and xenopsin expressed in the extraocular CPRs (**Figure 3K**).

**Figure 3.**
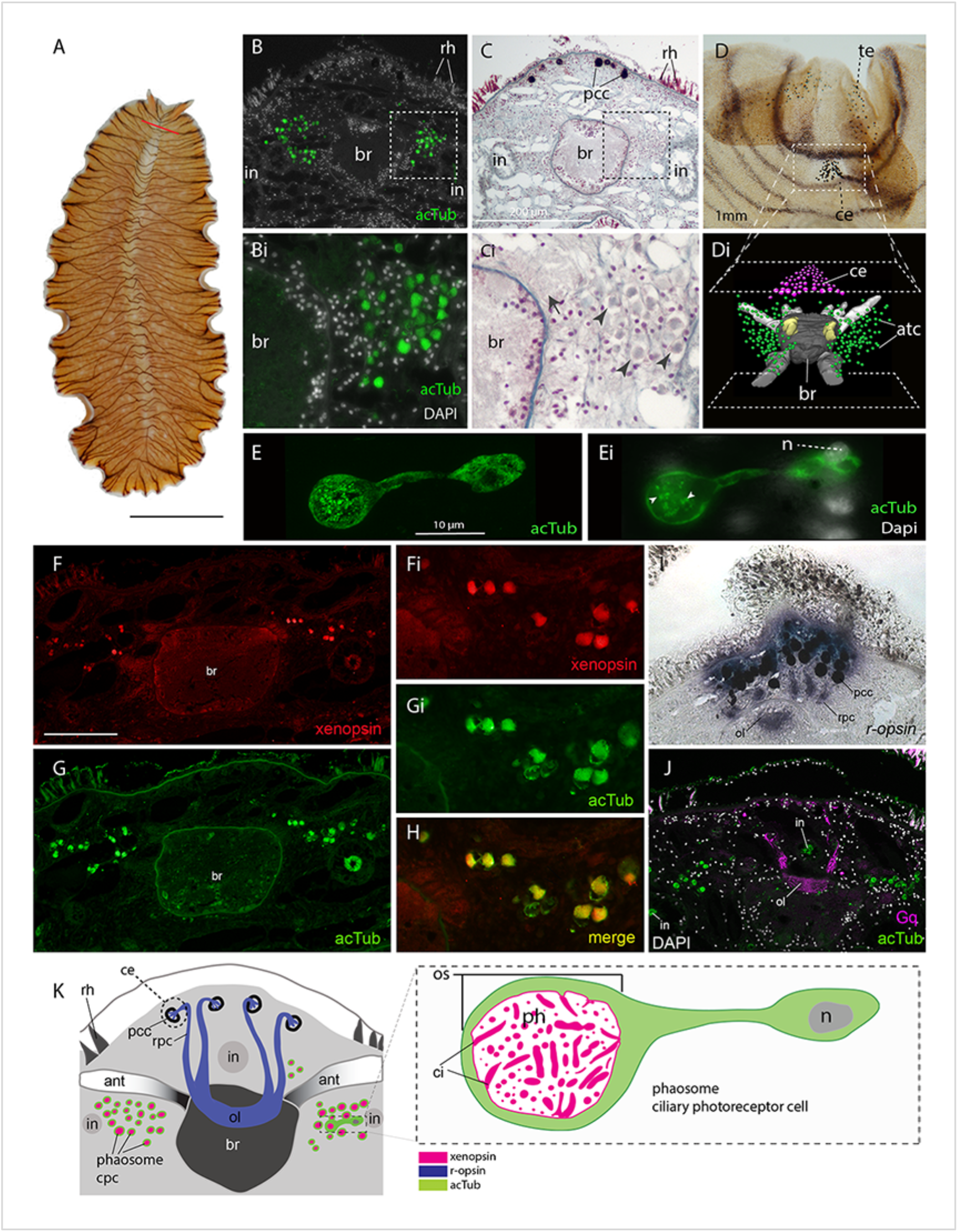
*Mc* xenopsin is co-expressed with acetylated tubulin in two dense clusters of extraocular, ciliary photoreceptors either side of the adult brain. A) Live adult (scale = 1cm), red line shows plane of cross section in B,C,G-K. B & C) Consecutive sections showing; B) two clusters of acetylated tubulin_+_ cells and; C) their distribution between the brain (*br*) and intestinal branches (*in*). Bi & Ci) Close up showing that these putative CPCs (arrowheads) are embedded in extracellular matrix in close proximity to main nerve tracts (arrow). Pigment cup cells (*pcc*), rhabdites (*rh*). D) Anterior end of adult showing pigmented eyespots above the brain (cerebral eyes, *ce*) and on the tentacles (tentacular eyes, *te*). Di) Schematised distribution of acetylated tubulin^+^ cells (*act*) and cerebral eyes on a micro-CT reconstruction of the brain and main anterior (white), posterior (grey), and optic (yellow) nerve tracts. E) Confocal projection of a putative CPR staining positive for acetylated tubulin and, Ei) an optical slice of the same cell showing position of nucleus (*n*) and cilia projecting into the intra-cellular vacuole (arrowheads). F-H) Co-localisation of acetylated tubulin and xenopsin suggests that these cells are photoreceptors (scale = 200µm);I) *r-opsin* expression in photoreceptor cells (rpc) that extend from the pigment cup cells (pcc) to the optic lobe (ol) of the brain; J) the position of the ciliary photoreceptors (labelled with acetylated tubulin) in relation to the rhabdomeric photoreceptors (labelled with Gq). K) Schematic of xen- and r-opsin expression in adult cross-section showing extraocular cpc and ocular rpc, diagram of ciliary photoreceptor showing outer segment (os) with xenopsin expression in phaosome containing cilia.

### 4. Serial SEM shows that extraocular CPRs in the adult have hundreds of cilia enclosed within a phaosome producing a large ciliary surface area

In order to characterize the morphology of the xenopsin^+^ cells in the adult we carried out serial SEM. These analyses show that these peri-cerebral cells house multiple, unmodified cilia in an intra-cellular vacuole and that they sit in close proximity to each other, in dense aggregations (**Figure 4A & B, and figure 4 video 1).** These cells are not associated with any pigmented supporting cells and, although there are unpigmented cells in close proximity (**Figure 4**), it seems as though the intra-cellular vacuole is completely enclosed by the cell itself, i.e. it can be considered a phaosome (**Figure 4 - video 2**).

The intracellular cavities/ phaosomes have diameters up to 23μm. The wall of the cavity is comparatively thin in certain areas (40 nm). The cytoplasm bordering the internal cavity contains mitochondria (**Figure 4C & D**). Within the phaosomal vacuole, cilia are anchored in the cytoplasmic layer bordering the intracellular lumen by basal bodies (**Figure 4D**) and each basal body gives rise to one cilium. Counting the basal bodies from the serial SEM of a single phaosome reveals at least 421 cilia projecting into the intracellular cavity (**Figure 4 - video 2**). The cilia are unbranched and emerge all around the diameter of the cavity (**Figure 4E, - video 3 and 4**), forming a tightly intertwined bundle (**Figure 4B**). They have an average diameter of 0.2μm and length of 7.2μm. This represents a total membrane surface area per phaosome of approximately 600μm^2^.

The cilia are generally orientated horizontally in relation to the dorso-ventral body axis and, in some of the phaosomes, the cilia appear to be arranged in a spirally coiled bundle (supplementary video 1). Near their bases, the axonemata show a 9×2+2 arrangement of microtubules. With increasing distance from the base, 9+2 singlets are encountered and then the nine-fold symmetry becomes disorganised and microtubular singlets are found (**Figure 4F; video 3**). As no ciliary rootlets were evident it is most likely that these cilia are non-motile; however, possible dynein arms (generally associated with motile cilia) were observed attached to the A-tubules near the bases of the cilia (Figure 5F). We observed these cells in live adults and although the cilia inside the phaosomes were visible, no cilia were seen moving, even in response to changes in illumination.

**Figure 4.**
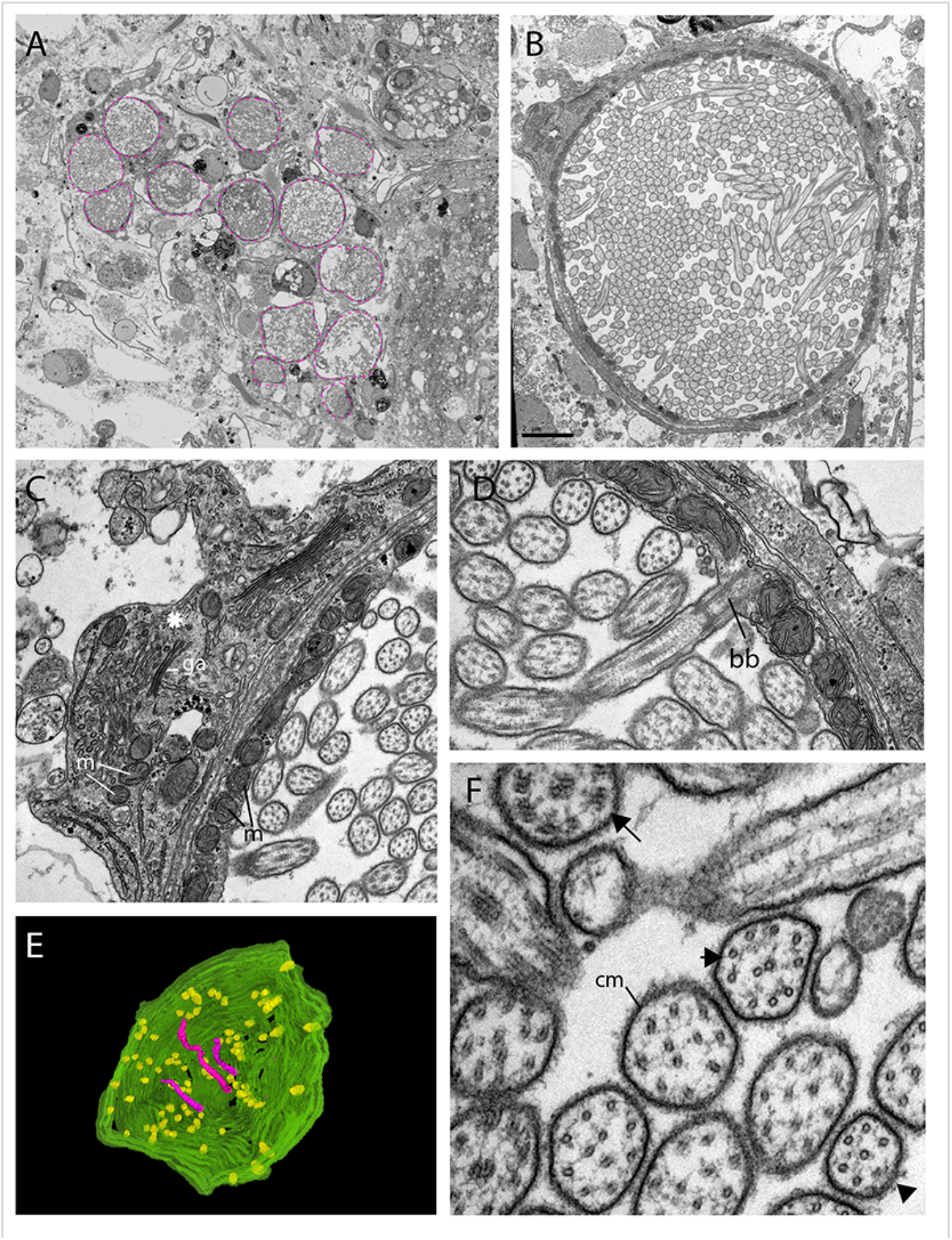
The morphology of the ciliary phaosomal photoreceptors in adult *Maritigrella crozieri.* A) A dense cluster of intra-cellular vacuoles (phaosomes – highlighted in magenta) filled with cilia which form the outer segment of the photoreceptor cell. B) Outer segment with multiple cilia in the phaosome. C) A possible unpigmented supporting cell (asterix) wrapping around the photoreceptor cell with mitochondria (m) and golgi apparatus (ga) in the cytoplasm. D) Ciliary axonemata (ax) are anchored in the cytoplasmic layer (cl) by basal bodies (bb). E) 3D reconstruction of the interior of a third of a phaosome, showing that the cilia are unbranched (pink) and the basal bodies (yellow) are distributed all around the phaosome. F) Cross sections of the ciliary axonemata show various arrangements of microtubules : 9×2+2 with dynein arms attached to the A-tubules (arrow), 9+2 singlets (double arrowheads), and singlets (triple arrowheads). This variation is related to the distance from the basal body (supplementary video 3). Scale in (B) = 2µm, (cm) ciliary membrane.

**Figure 4 – video 1:**
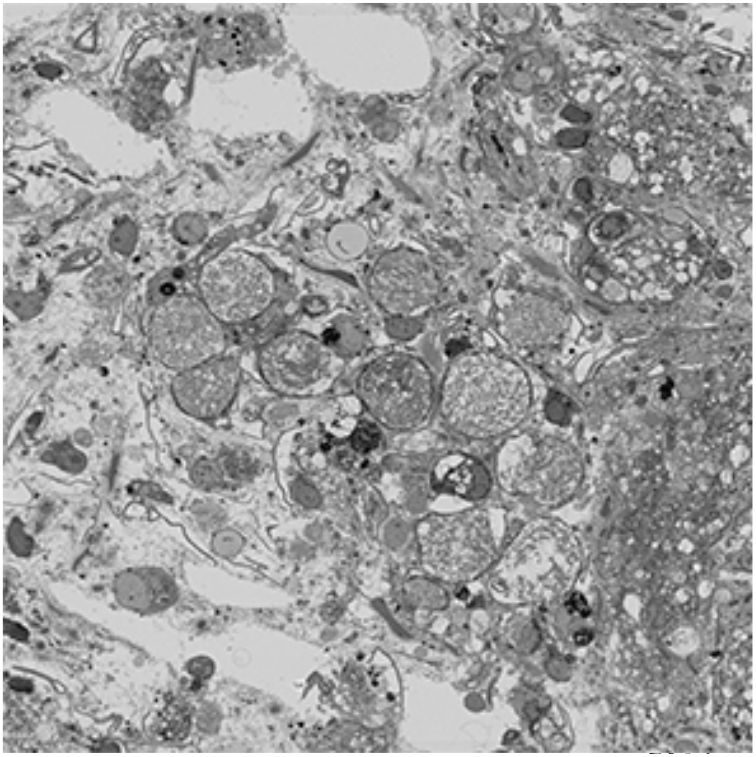
Serial SEM images (101 × 500nm sections = 50.5μm total thickness) showing a cluster of ciliary phaosomes (intra-cellular vacuoles housing multiple cilia) that form the outer segment of the extraocular CPR cells in adult *Maritigrella*. This cluster is located to the left of the brain.

**Figure 4 – video 2:**
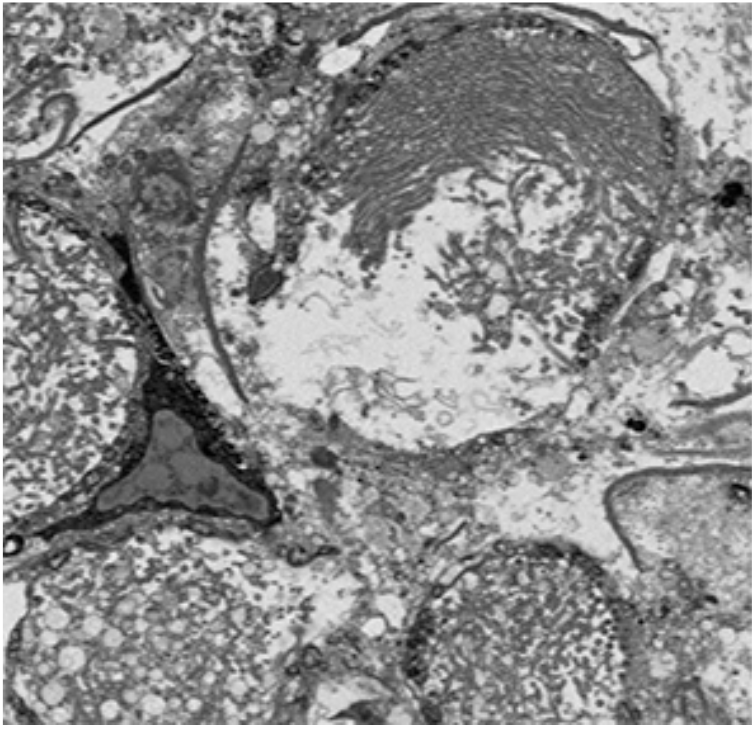
Serial SEM images (47 × 400nm sections = 18.8μm total thickness) showing a complete phaosome. At least 421 basal bodies were counted projecting cilia into the intra-cellular space that is formed by the cell itself. Note that in a nearby phaosome the cilia appear to be arranged in a spirally coiled bundle.

**Figure 4 – video 3:**
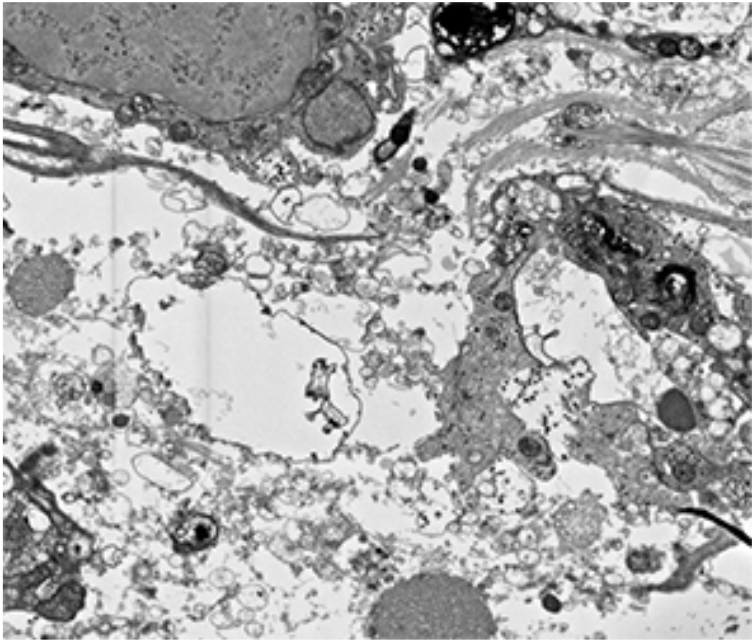
Serial SEM images (72 × 100nm sections = 7.2μm total thickness) showing a third of a phaosome.

**Figure 4 – video 4:**
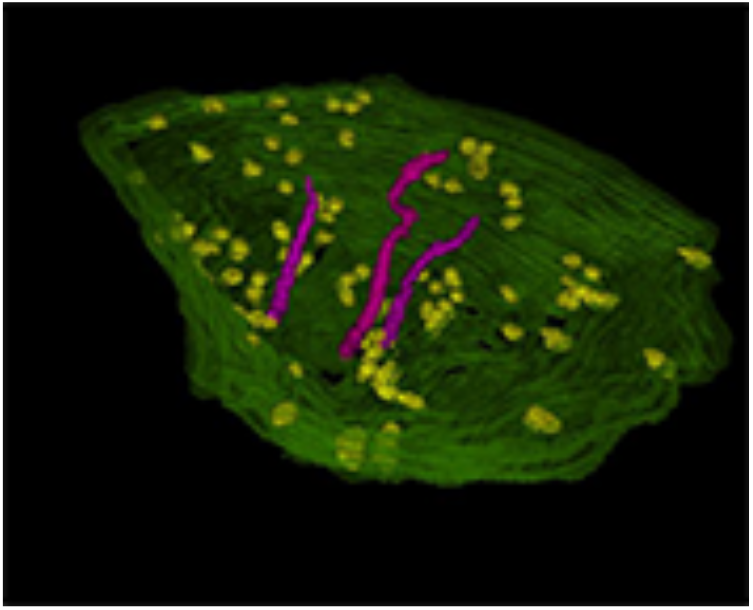
A 3D reconstruction of series in video 3 shows that the cilia are unbranched and emerge all around the diameter of the cavity.

**Figure 5.**
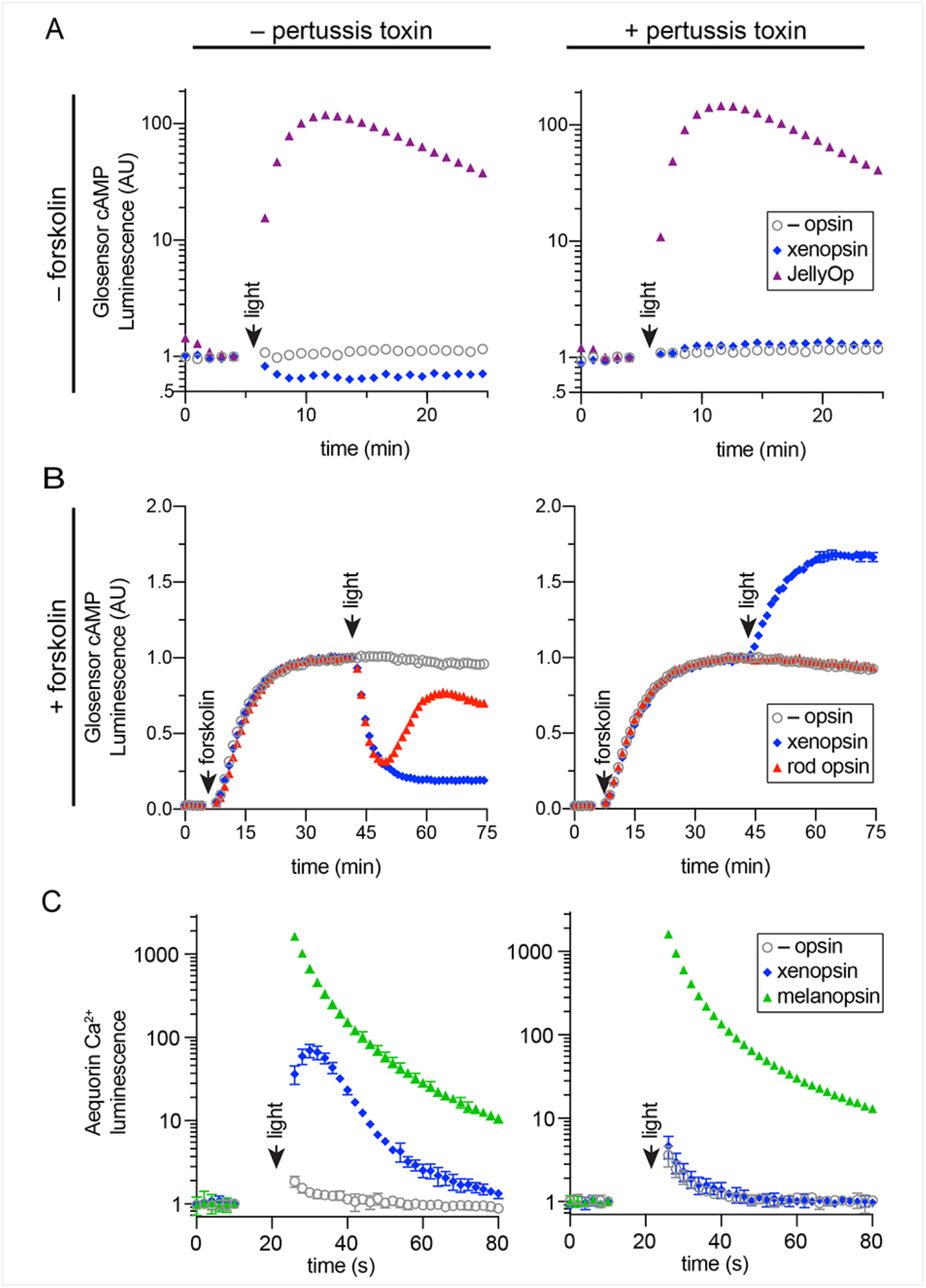
In human cells *Maritigrella crozieri* xenopsin forms a functional photopigment that predominantly couples to Gi/o pathways. a,b) HEK293T cells were transfected with Glo22F and indicated opsins, +/-PTX, and exposed to light. In b, cells were treated with 2uM forskolin prior to light flash. c) HEK293T cells were transfected with mtAequorin and indicated opsins, +/-PTX, and exposed to light. Plots show mean luminescence of technical replicates (from one representative of three biological replicates) normalized to the pre-flash timepoint, +/-SEM. Error bars smaller than symbols are not shown. n=3 technical replicates in a,b; n=4 technical replicates in c. The other biological replicates are shown in **Supplementary figure 4**.

### 5. Maritigrella xenopsin forms a photopigment capable of sustained G α _i/o_ signalling

To determine whether *Maritigrella* xenopsin functions as a photopigment, and to explore which class(es) of G protein it can couple to, we assessed its ability to modulate levels of the second messenger molecules cyclic AMP (cAMP) and calcium (Ca^2+^) in response to light when heterologously expressed in HEK293 cells (Bailes and Lucas, 2013; Koyanagi et al., 2013). Changes in cAMP or Ca^2+^ levels are characteristic of opsin coupling to the three major classes of G alpha protein: Gs, Gi/o, and Gq. To a first approximation, Gs coupling causes an *increase* in cAMP; Gi/o coupling causes a *decrease* in cAMP; and Gq coupling causes an increase in cytoplasmic Ca^2+^. We compared *Maritigrella* xenopsin to three positive control opsins: the cnidopsin JellyOp, human rod opsin, and human melanopsin, which are known to potently and selectively couple to Gs, Gi/o, and Gq, respectively (Bailes et al., 2012; Bailes and Lucas, 2013). In all experiments, we used a flash of 10^15 photons of 470 nm light as the stimulus. Second messenger levels were monitored in real time using the bioluminescent reporter proteins Glosensor cAMP 22F (Glosensor) for cAMP and Aequorin localized to the cytoplasmic surface of the mitochondria (mtAequorin) for cytoplasmic Ca^2+^.

To assay for Gs coupling (**Figure 5A**), cells were transfected with either xenopsin or JellyOp, and Glosensor and exposed to light. As expected, JellyOp induced a >100-fold increase over baseline in Glosensor luminescence, and no response was observed in negative control cells without opsin. By contrast, xenopsin induced a ~40% decrease in Glosensor signal in response to light, suggesting Gi/o coupling. The addition of pertussis toxin (an enzyme that inactivates Gi/o) blocked the decrease in cAMP in xenopsin-expressing cells indicating that it was caused by coupling to Gi/o. We also observed that xenopsin drove a small 0.2 fold increase in luminescence during pertussis toxin treatment, suggesting that xenopsin may also weakly couple to Gs. JellyOp signaling was not affected by the addition of pertussis toxin, which does not interfere with Gs.

To better assay for Gi coupling, we used the drug forskolin to artificially elevate cAMP by activating adenylyl cyclase prior to light flash. HEK293 have low basal levels of cAMP, so forskolin treatment increases the magnitude of cAMP suppression that is possible to achieve with a Gi/o coupled opsin. Cells were transfected with Glosensor and xenopsin or rod opsin, with or without pertussis toxin pretreatment. After measuring basal cAMP levels, forskolin was added, which induced a strong increase in Glosensor signal that stabilized after ~40 minutes. At this point, cells were exposed to light. Both rod opsin and xenopsin suppressed cAMP to below 50% of pre-flash levels, whereas no decrease was seen in cells without opsin. As expected, the ability of rod opsin and xenopsin to suppress cAMP was entirely blocked by pertussis toxin, confirming the role of Gi/o in this activity (**Figure 5B**).

Although xenopsin and rod opsin both suppress cAMP by coupling to Gi/o, there are intriguing differences in the responses they induce. Rod opsin produces a transient decrease in cAMP, which reaches a minimum at 5 minutes post-flash and returns to the level of control cells by 20 minutes. In contrast, xenopsin appears to irreversibly suppress cAMP in this system (**Figure 5B**). The long lifetime of the xenopsin response indicates that its signaling active state is very stable in this system, and suggests that the signal termination mechanisms (e.g. GPCR kinases and beta-arrestins) present in HEK293 cells may not be suitable for xenopsin. Surprisingly, we also observed cAMP increase in pertussis toxin treated cells in the presence of forskolin (**Figure 5B**). This light response implies that although xenopsin can signal via Gi/o it must also have some additional signaling capacity that is capable of increasing cAMP. Our combined Glosensor data suggest that xenopsin may be a promiscuous opsin that is capable of increasing cytosolic cAMP through an as yet undefined pertussis toxin-insensitive pathway, but that Gi/o coupling predominates unless it is artificially inactivated.

Finally, we tested the ability of xenopsin to modulate cytoplasmic Ca^2+^ release, in comparison to melanopsin, with or without pertussis toxin pretreatment. It is known that Gi/o activation can trigger cytoplasmic Ca^2+^ release. In cells transfected with melanopsin, light induces a >1000-fold increase in cytoplasmic Ca^2+^, which is not affected by pertussis toxin. Xenopsin triggers a smaller ~100-fold increase in cytoplasmic Ca^2+^, importantly, this response is almost entirely abolished by pertussis toxin (**Figure 5C**). Because we did not detect pertussis-toxin independent Ca^2+^ signalling activity, we conclude that xenopsin is not capable of coupling to Gq. Taken together these results show that in human cells, xenopsin forms a functional opsin that predominantly suppresses cAMP in response to light, by coupling to Gi/o pathways, and that it may have a minor capacity to couple to Gs or another undefined pathway.

## Discussion

Our phylogenetic analyses of opsin genes across the Metazoa supports the emerging consensus that xenopsins and c-opsins diverged prior to the bilaterian common ancestor. As both opsin types are found in various protostomes, this suggests that the two opsins co-existed in the protostome stem lineage. The known taxonomic distribution of xenopsins and c-opsins is strange in this context in that no species is known in which both opsins are present. Our survey of flatworm opsins has revealed another instance in which only one of the two related xenopsin/c-opsins types is found.We find that the same is true of a chaetognath and a bryozoan. Why different groups have retained one opsin rather than the other is unknown and whether any species exist with both opsins will require broader sampling.

One possibility to explain this distribution is that, while clearly separate clades with distinct and conserved genes structures unique to each type of opsin, the two opsins are, nevertheless, largely equivalent. In addition to their relatively close phylogenetic relationship, it is obvious that both opsins are expressed in cells with expanded ciliary membranes. We have now gone beyond comparative analysis of amino acid sequences by demonstrating that xenopsins, like c-opsins, are capable of activating a Gαi signal transduction cascade in living cells. We have shown further similarities in the shared possession of the VxPx motif which, in c-opsins, targets the protein to the ciliary membrane. We have also shown that this motif extends to some cnidopsins, which cluster together with xenopsins and c-opsins in phylogenetic analyses (Fig.1).

Our analyses also support the idea that xenopsins themselves duplicated as suggested by the presence of two clades of xenopsin sequences from the polyclad flatworms (Vöcking et al., 2017). Xenopsins from clade A have the signatures to be photopigments, while those in clade B although they have lysine in transmembrane domain VII to bind to the chromophore, lack the NxQ and VxPx motifs for Gα _i_-protein binding and transport to cilia. This might suggest that they are photoisomerases supporting the xenopsin photopigments of clade A by recycling the chromophore.

We have provided the first evidence that a xenopsin, *Mc* xenopsin, forms a functional photopigment. When heterologously expressed in HEK293 cells, and reconstituted with 9-*cis* retinal, *Mc* xenopsin elicits a light-dependent decrease in cAMP that is blocked by pertussis toxin, indicating that it can signal via a Gα _i_ signal transduction cascade. *Mc* xenopsin also drove a transient increase in intracellular Ca^2+^ of the type elicited by Gq coupled opsins, but as this response was blocked by pertussis toxin it likely reflects crosstalk with the Gαi signaling pathway. Nevertheless, our data do indicate that *Mc* xenopsin has promiscuous signaling. Thus, when Gα _i_ is inactivated by pertussis toxin, *Mc* xenopsin acts to increase cAMP in response to light, revealing additional coupling to a pertussis toxin insensitive pathway. While increases in cAMP can be elicited by Gs driven increases in adenylyl cyclase activity, we do not think that this is the case for *Mc* xenopsin, as the cAMP response was strongly potentiated by forskolin (which also activates adenylyl cyclase).The alternative is that *Mc* xenopsin couples to some as yet undefined, pertussis toxin insensitive cascade as proposed previously for other Gi/o coupled invertebrate opsins, such as scallop opsin 1 (Ballister et al., 2018). Nonetheless, in the case of xenopsin, cAMP suppression through Gα _i_ coupling is the dominant effect in unperturbed cells indicating that this is a dominant signaling mode.

Compared to human rod opsin, *Mc* xenopsin was effectively irreversibly activated by blue light in this assay; there was no change in its signaling over tens of minutes. Although this may reflect an incompatibility with human GPCR kinases and arrestins, it also implies that the activated state of the opsin is thermally stable, and may be bistable, with a thermally stable signal state that can be converted back to inactive dark state by subsequent light absorption (Tsukamoto and Terakita, 2010). Bistable opsins, such as lamprey parapinopsin, are known to exhibit prolonged cAMP suppression in response to blue light in live cell assays of Gi activation (Kawano-Yamashita et al., 2015), similar to the responses observed with xenopsin. Several aspects of xenopsin signaling must be explored further. What are the relevant second messengers and signaling kinetics in its native phaosome context, and which G alpha proteins are expressed there? What are the spectral sensitivity, quantum efficiency, and cofactor requirements of xenopsin? These questions may be addressed by a combination of *in vivo* electrophysiology and *in vitro* cellular assays. The results presented here provide a foundation for further detailed functional investigation of this clade of opsins.

We have shown that xenopsin is expressed in two functionally and morphologically different types of CPRs at different developmental stages in *Maritigrella crozieri*; ocular expression in the larval stage but an extraocular, non-visual role in the adult. Although xenopsin has been described as a visual opsin (Vöcking et al., 2017) due to expression in putative eyes of the larval stages of a brachiopod (Passamaneck et al., 2011) and chiton (Vöcking et al., 2017), it was also found to be expressed in extraocular photoreceptors of the chiton larva (Vöcking et al., 2017). Xenopsin expression in both ocular and extraocular photoreceptors, either simultaneously (e.g. in the chiton larva (Vöcking et al., 2017)) or sequentially over development (e.g. in the larval eyespot and adult extraocular PRCs of *Maritigrella*), is a feature common to most other opsin clades (Porter et al., 2012; Cronin and Porter, 2014; Sprecher and Desplan 2008).

The adult, extraocular, non-visual xenopsin^+^ cells of *Maritigrella* resemble the extraocular CPRs of the marine annelid *Platynereis.* In the annelid these are located in the brain and express a UV-sensitive ciliary opsin (Arendt et al., 2004; Tsukamoto et al., 2017). These cells control circadian behaviours via melatonin production (Tosches et al., 2014) and mediate downward swimming in response to non-directional UV light in the larval stage (Verasztó et al., 2018). The *Platynereis* c-opsin binds to exogenous all-*trans*-retinal, which is particularly important for opsins expressed outside of the eyes (Tsukamoto et al., 2017) where sophisticated multi-enzyme systems producing a thermally unstable 11-*cis*-retinal isomer probably don’t exist (Yau and Hardie, 2009). Although *Mc* xenopsin was able form a photopigment with 9-*cis*-retinal it would be interesting to examine retinal isomer preferences.

The unique morphology of the flatworm extraocular ciliary photoreceptor outer segment seems to be a flatworm novelty (Sopott-Ehlers et al., 2001). A comparison of the morphology of *Maritigrella* phaosomal CPRs with those of other Metazoa shows two lines of evidence that support this idea (Fig.6). Firstly, while most metazoan CPRs project their ciliary membranes outside of the cell (like jawed vertebrate cones), *Maritigrella* extraocular CPRs enclose their ciliary membranes within their plasma membrane, like gnathostome rods and a few examples from other invertebrates (Fig. 6). The patchy phylogenetic distribution of cells enclosing their ciliary processes suggests these have probably evolved independently in these taxa. The enclosure of ciliary processes in gnathostome rods and invertebrate phaosomes also proceeds in different ways (evagination of the ciliary plasma membrane leading to enclosed pinched off discs in rods (Steinberg et al., 1980), versus the presumed invagination of the apical cell membrane during development in phaosomes (Purschke, 2003; Purschke et al., 2006)) (Fig. 6). Secondly, *Maritigrella* CPRs increase their surface area by increasing the number of cilia rather than modifying the ciliary membranes into discs or lamellae (Fig.6). The sheer number of cilia housed within the phaosome is striking, and may be a unique feature of *Maritigrella* and other flatworm CPRs. A few examples of unmodified cilia in complete or open phaosomes have been recorded in other lophotrochozoans (Hessling and Pursche, 2000; Wollacott and Zimmer, 1972), but most other phaosomal CPRs have modified cilia (flattened and whorled) (Clement and Wurdak, 1984; Boyle, 1956; Horridge, 1964). The function of enclosing the ciliary membranes is not known, even in rods, but suggestions include increased efficiency in the transport of photopigments, the renewal of the outer segments, and of signalling or protein function with ion channels on the plasma membrane separated from opsins and other transduction proteins on the ciliary membrane (Morshedian and Fain, 2015; 2017). Our study - the first to document the expression of an opsin within an invertebrate ciliary phaosome - will facilitate future comparative studies between phaosomes and rods to understand the function and evolution of this morphology.

**Figure 6.**
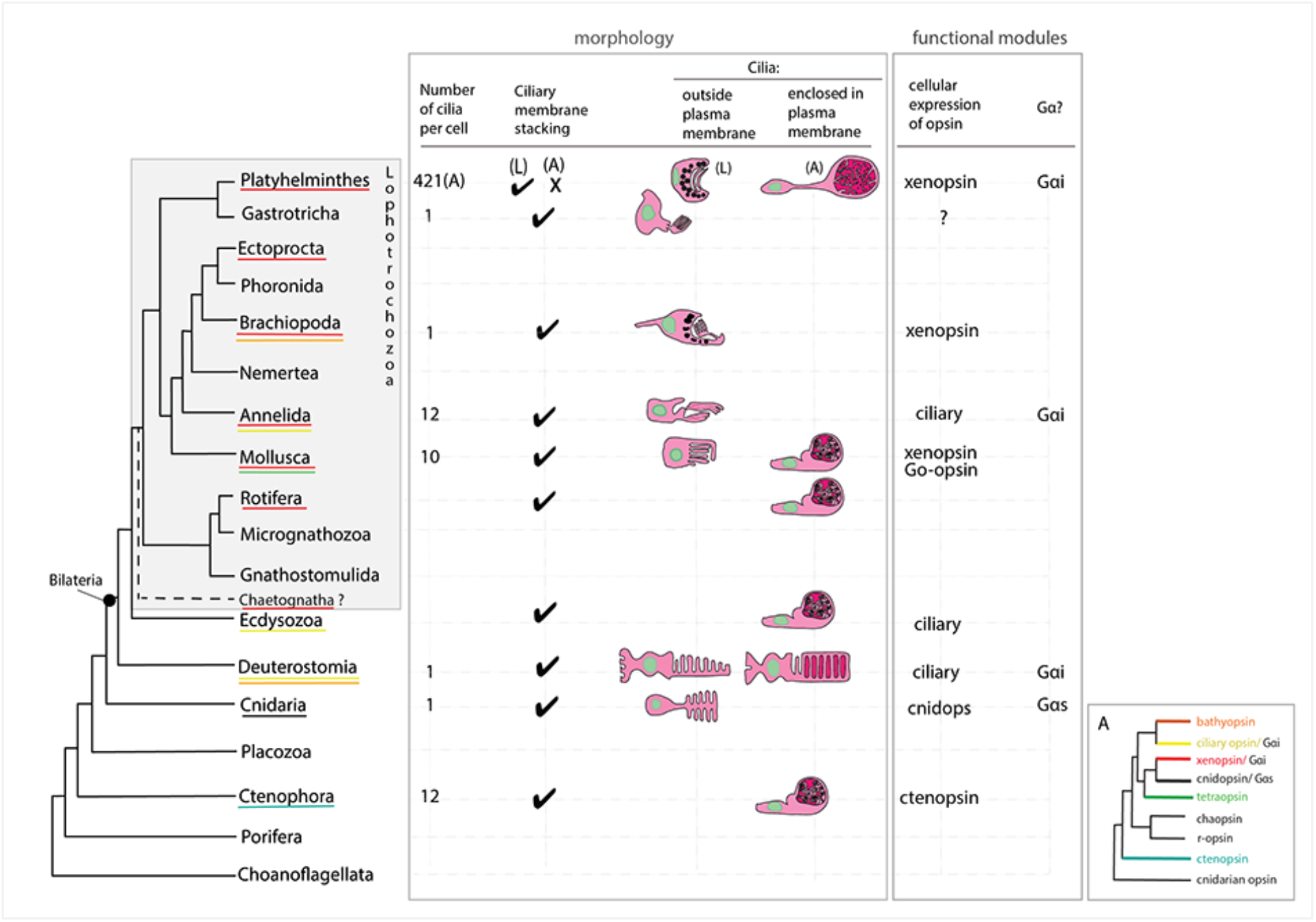
An overview of metazoan ciliary photoreceptor morphology, opsin expression and G-protein coupling (determined from cellular assays), highlighting the distinct morphology of the ciliary phaosomes in flatworms and possible convergent evolution of enclosed ciliary membranes in invertebrate phaosomes and jawed vertebrate rods. The coloured lines under the phylum names represents the presence of the opsin sub-type in the taxonomic group; note the higher prevalence of xenopsins over ciliary opsins in Lophotrochozoa. (L) = larval photoreceptor, (A) = adult photoreceptor. Box A shows the opsin relationships according to our phylogeny and the known Gα-binding of opsins expressed in ciliary photoreceptors.

Our morphological analyses show that the phaosomal CPRs of *Maritigrella* adults are not associated with pigment cup cells and are therefore extraocular and could detect light from all directions.Animals use non-directional light to set circadian cycles, monitor UV levels and photoperiodism, gauge depth, and detect a predator’s shadow (Nilsson, 2009). However, cells that only signal slow changes of ambient light intensity can work without membrane specializations (Nilsson 2004), such as intrinsically photosensitive retinal ganglion cells (Hattar et al., 2002). The density of CPRs in *Maritigrella*, and the number of cilia in each phaosome, increases the surface area to allow higher concentrations of photopigment, which would suggest high sensitivity to light. The ciliary surface area of a phaosome is approximately 4 times smaller than the total disc membrane surface of rat rods (Mayhew and Astle, 1997) but 2 times larger than that of the brain CPRs of the annelid *Platynereis* (Verasztó et al., 2018). High sensitivity is normally associated with a visual role (Nilsson, 2004) but as all extraocular photoreceptors are, by definition, non-visual (Cronin and Johnsen, 2016) then their membrane elaborations could indicate that these photoreceptors function to detect intermediate to fast changes of non-directional light associated with changes in depth or a predator’s shadow, rather than (or as well as) the slower changes of light over 24 hours or seasons. If *Mc* xenopsin in its native cell shows the same slow responses that we show here then perhaps these cells function to detect and amplify low levels of light, like vertebrate rods, and could be involved in detecting moonlight.

Marine animals such as *Maritigrella crozieri* could use light in many ways. In addition to light as a visual stimulus, light intensity and wavelength can provide clues as to the time of day, the season, the state of the tide and water depth. Light will also be used differently by a minute swimming larva and a large crawling adult worm. We have characterised a new type of opsin-expressing cell in a flatworm and demonstrated that the opsin is capable of photosensitivity and phototransduction. This adds to the increasing diversity of animal photoreceptors and phototransduction pathways being discovered as more species are studied.

## Methods

### Identification of ciliary-type and rhabdomeric opsin sequences

*Maritigrella crozieri* ciliary-type and rhabdomeric opsins were identified by reciprocal best match BLAST searches on a mixed stage embryonic and larval transcriptome [Lapraz et al., 2013]. Four *Schmidtea mediterranea* opsin sequences [Zamanian et al., 2011 – Additional File 7] along with the *Maritigrella* sequences were used, with Mollusc opsins [Ramirez et al., 2016], as query sequences for BLAST searches against assembled transcriptomes and genomes for 30 other flatworm species [Egger et al., 2015, Laumer et al., 2015], a chaetognath (*Pterosagitta draco*) and a bryozoan (*Bugula neritina*). Our search allowed the identification of some already published flatworm sequences [Vöcking et al., 2017] (DAB27256.1, DAB27257.1, DAB27258.1, DAB27259.1, DAB27253.1, DAB27254.1 and DAB27255.1). These published accession numbers and sequences were used in our analysis.

### Phylogenetic analysis

Flatworm, bryozoan and chaetognath best hit sequences were added to a subset of the Ramirez *et al.* (2016) metazoan opsin sequences dataset. The subset was obtained by first reducing redundancy of the original dataset using an 80% identity threshold with CD-HIT (Li and Godzip, 2006), then by discarding sequences which, on an alignment, did not fully cover the region found between first and last transmembrane region. Finally, when multiple sequences belonging to the same taxonomic clade or class were found, only the two or three most complete representative sequences where kept. Additional opsin sequences were added to the dataset: human melanopsin, human rhodopsin and *Carybdea rastonii* opsin (called JellyOp in this study), *Xenopus laevis* OPN4B (covering a taxonomic gap), *Acromegalomma interruptum* InvC-opsin and *Spirobranchus corniculatus* InvC-opsin (kindly provided by Dr. Michael Bok, (Bok et al., 2017)), *Owenia_fusiformis* Xenopsin1, 2 and 3 (Vöcking et al 2017), *Platynereis dumerilii* TMT1 (http://genomewiki.ucsc.edu/index.php/Opsin_evolution), as well as additional non-opsin outgroup sequences (Prostaglandin E2 receptor and Melatonin receptor sequences) resulting in a first dataset (Dataset 1) of 213 sequences. In order to evaluate their influence on the tree topology, sequences forming small monophyletic groups (Bathyopsin, Chaopsin, ctenophore and cnidarian early branching opsins in Ramirez *et al.* opsin phylogeny (Ramirez et al., 2016)) were removed from our initial dataset (Dataset 2 – 196 sequences).

For both datasets, sequences were aligned with MAFFT (Katoh and Standley, 2013) webserver (https://mafft.cbrc.jp/alignment/server/) using the L-INS-I option. Portions of the alignment with fewer than 6 represented positions were trimmed from the alignment using trimAl v1.2rev57 (Capella-Gutiérrez et al., 2009), then the alignment was manually trimmed to remove positions before first aligned methionine and after the last aligned block.

For both datasets, Maximum-Likelihood phylogenetic reconstruction of the trimmed alignment was conducted using both: IQ-TREE webserver (http://iqtree.cibiv.univie.ac.at/) (Trifinopoulos et al., 2016) with a LG+R9+F substitution model, and with 1000 Ultrafast bootstrap replications as well as SH-aLRT (1000 replicates) and approximate aBayes single Branch testing, or with RAxML v.8.2.9 (Stamatakis 2014) on the Cipres webserver (www.phylo.org/portal2/) (Miller et al., 2010) with a GAMMA-LG-F substitution model and 100 rapid boostrap replicates. FigTree v1.4.3 (tree.bio.ed.ac.uk/software/figtree/) was used for tree visualisation. Accession numbers of the sequences used in the phylogenetic analysis are available in the supplementary table 1.

### The morphology, and opsin expression, of Maritigrella crozieri ciliary photoreceptors

#### Animal collection, fixation and sectioning

Adult *Maritigrella crozieri* were collected from the Florida Keys (Rawlinson, 2010; Lapraz et al., 2013). They were fixed in a Petri-dish containing frozen 4% paraformaldehyde (diluted in phosphate buffered saline [PBS]) overnight at 4°C, rinsed in PBS (3 × 5 minutes, 5 × 1 hour washes) at room temperature and dehydrated in a step-wise ethanol series for histology and immunofluorescence, and in a methanol series for mRNA *in situ* hybridisation. For histology, heads of adult worms (from the pharynx anteriorward) were dissected, cleared in histosol (National Diagnostics), and embedded in paraffin. Paraffin blocks were sectioned at 8-12µm using a Leica (RM2125 RTF) microtome. Larval stages were fixed in 4% PFA in PBS for 20 minutes at room temperature, rinsed in PBS for five × 30 minute washes and stored in 1% PBS-azide at 4°C for immunofluorescence, or dehydrated into 100% methanol and stored at −20°C for mRNA *in situ* hybridization.

#### Histology and immunohistochemistry

For adult stages, consecutive sections were used to compare histology and immunofluorescence. For histological analysis, sections were stained with Masson’s trichrome (MTC) [Witten and Hall, 2003]. For immunostaining of paraffin sections, slides were dewaxed in Histosol (2×5 min), then rehydrated through a descending ethanol series into PBS + 0.1% Triton (PBT, 2×5 min). Slides were blocked with 10% heat-inactivated sheep serum in PBT for 1 hour at room temperature in a humidified chamber. Primary antibodies (see below) were diluted in block (10% heat-inactivated sheep serum IN PBT) and applied to the slide, covered with parafilm, and incubated at 4°C for 48 hours. Slides were then rinsed in PBT (3×10min). Secondary antibodies diluted in block solution were then applied to each slide, and slides were covered with parafilm and incubated in a humidified chamber, in the dark, at room temperature for 2 hours. Slides were rinsed in PBT 3 × 10 minutes, and then 4 × 1 hour prior to counterstaining with the nuclear marker 4’, 6-diamidino-2-phenylindole (DAPI) (1 ng/ml) and mounting in Fluoromount G (Southern Biotech, Birmingham, AL). Immunostaining of larval stages was performed according to Rawlinson (2010).

Primary antibodies used were: anti-acetylated tubulin (Sigma) diluted at 1:500, a polyclonal antibody directed against the C-terminal extremity of the *Maritigrella* xenopsin protein sequence (GASAVSPQNGEESC; generated by Genscript, Piscataway, NJ, USA) diluted at 1:50, and anti-Gq/11α (C-19, Santa Cruz Biotechnology, Santa Cruz, CA, USA) diluted at 1:300. Imaging of immunofluorescence on paraffin sections and larval wholemounts was carried out using an epi-fluorescent microscope and a confocal laser scanning microscope, additional images on larvae were taken with an OpenSPIM (Girstmair et al., 2016). For the 3D rendering of the larva a multi-view stack was produced by capturing several angels of the specimen and using Fiji’s bead based registration software and multi-view deconvolution plugins (Preibisch 2010, Preibisch 2014).

#### mRNA in situ hybridization

To analyse the expression of *Maritigrella crozieri r-opsin*, we performed mRNA *in situ* hybridization using a riboprobe generated against the *r-opsin* sequence identified above. A 523bp fragment of *M. crozieri r-opsin* was PCR amplified using the following primers: *r-opsin*-fw TCCCTGTCCTTTTCGCCAAA, *r-opsin*-rv TATTACAACGGCCCCCAACC. The fragment was cloned using the pGEM-T easy vector system, and a DIG-labelled antisense probe was transcribed according to the manufacturer’s protocol. mRNA *in situ* hybridization on paraffin sections of adult tissue was carried out according to O’Neill et al. [2007]. Upon completion of the colour reaction, slides were coverslipped with Fluoromount G. Wholemount mRNA *in situ* hybridization on larvae was carried out according to the *Capitella tellata* protocol of Seaver and Kaneshige (2006). Following termination of the colour reaction, specimens were cleared and stored in 80% glycerol, 20% 5× PBS. Both adult and larval mRNA *in situ* hybridization experiments were imaged on a Zeiss Axioscope.

#### TEM and serial SEM

Adult *Maritigrella crozieri* heads were dissected and immediately placed in ice-cold, freshly prepared 3% glutaraldehyde overnight at 4 °C. The tissue was rinsed seven times in 0.1M sodium phosphate buffer, pH 7.2, then placed in 1 % osmium tetroxide (in the same buffer) for 1 hour at 4°C. Samples were then rinsed twice with ice-cold distilled water and dehydrated in an ethanol series (50 %, 75 %, once each for 15 min; 95 %, 100 % twice each for 15 min), culminating in two changes of propylene oxide with a waiting period of 15min after each change. The samples were then placed in Epon mixture/propylene oxide (1:1) for 45min at room temperature (22–25 °C).Finally, samples were transferred from vials into fresh Epon mixture in molds and polymerized in an oven at 60 °C for 72 h.

For TEM, sections of 60-70nm thickness were cut with a diamond knife on a Reichert Ultracut E ultramicrotome. After their collection on formvar film coated mesh grids, the sections were counterstained with lead citrate. The ultrathin sections were analysed using a Jeol-1010 electron microscope at 80 kV mounted with a Gatan Orius camera system.

For serial SEM the samples were shaped to an ~1 × 4 mm rectangular face using a diamond trimming tool. The block was mounted in a microtome (Leica EM UC7, Buffalo Grove, IL) and thin sections, 50-100 nm in thickness, were cut with a diamond knife. The methods are described in detail in Terasaki et al. (2013) but in brief the sections were collected on kapton tape with the ATUM tape collection device, the tape containing the sections was cut into strips, mounted on 4 inch silicon wafers and then carbon coated. The sections were imaged using a field emission scanning EM (Zeiss Sigma FE-SEM, Peabody) in backscatter mode (10 keV electrons, ~5 nA beam current). The images were aligned using the Linear Alignment with SIFT algorithm and reconstructed using TrakEM2, both in FIJI Image J (Cardona et al., 2012). To estimate the sensory membrane surface area of a the phaosomal CPRs we counted the number of basal bodies (in two complete phaosomes), and calculated the average diameter and total length of 3 cilia per phaosome.

#### Micro-CT analysis

One adult *Maritigrella crozieri* was fixed in 4% PFA, rinsed in PBS and dehydrated into methanol, as described above. It was then stained in 1% (w/v) phosphotungstic acid (Sigma 221856) in methanol for 7 days, with the solution changed every other day. The animal was rinsed in methanol, mounted in an eppendorf tube between two pads of methanol-soaked tissue paper, and scanned on a Nikon XTH225 ST at the Cambridge Biotomography Centre (Department of Zoology, University of Cambridge). The brain area was segmented using Mimics software (Materialise, Leuven, Belgium).

### G-protein selection of Maritigrella crozieri xenopsin

We followed the methods for the secondary messenger assays as described in detail in Bailes and Lucas (2013). In brief, a mammalian expression vector was constructed using pcDNA3.1 (Invitrogen) and the open reading frame of *Maritigrella* xenopsin with the stop codon replaced by a 6 base linker and 28 bases that code for the 1D4 epitope from bovine Rh1 opsin. Expression vectors for the positive controls (Gs – Jellyfish opsin [JellyOp]; G_i_ – human rhodopsin [Rh1]; G_q_ – human melanopsin [Opn4]) were constructed in the same way (Bailes and Lucas 2013). Opsin-expressing plasmids were omitted from transfection in the negative controls. To make an expression plasmid for a luminescent cAMP reporter, the region for the Glosensor cAMP biosensor was excised from pGlosensor 22 (Promega) and ligated into linearized pcDNA5/FRT/TO. All restriction enzymes were from New England Biolabs (NEB). A luminescent calcium reporter was synthesized using the photoprotein aequorin from *Aequorea victoria* mtAeq (Inoue *et al*. 1985; Bailes and Lucas 2013).

#### Reporter and opsin transfection for light response assays

~6×10^4^HEK293 cells were plated per well in a 96 well plate 24hours prior to transfection in DMEM/10% FCS. Transfections were carried out using Lipofectamine 2000 (Invitrogen). Reporter and opsin-expressing plasmids were co-transfected at 500ng each and incubated for 4-6h at 37°C. DMEM/10% FCS + 10uM 9-*cis* retinal (Sigma) was then replaced and cells were left overnight at 37°C. All steps following initial transfection were carried out in dim red light only.

#### Luminescent second messenger assays

We tested three biological replicates per treatment, with each biological replicate consisting of an average of three technical replicates (for cAMP assays) or four technical replicates (for Ca2+ assays).

#### cAMP increases: Gs

For measurements of cAMP increases as an indication of G_s_ activity, wells of cells were transfected with pcDNA/FRT/TO Glo22F and opsin. Following transfection and overnight incubation, media was replaced with L-15 medium, without phenol red (Invitrogen), 10% FCS with 2mM beetle luciferin (Promega) for 1-2 hours at room temperature. Luminescence of the cells was measured with a Fluostar optima plate reader (BMG Labtech). After 6 minutes, cells were exposed to a flash of 470nm light (10^15^ photons) followed by a recovery period where relative luminescence units (RLU) were recorded every minute for up to 25 minutes.

#### cAMP decreases: Gi/o

Decreases in cAMP are difficult to measure from baseline cAMP reporter luminescence and so cells were treated with 2uM forskolin to artificially raise cAMP levels at 6 minutes. Luminescence was measured before and after the forskolin addition until the increase in luminescence plateaued. Cells were then flashed with 470nm light (as above) and luminescence measured for up to 45mins.

#### Ca2+ increases: Gq/11

Cells transfected with pcDNA5/FRT/TO mtAeq and opsins were incubated with 10uM Coelenterazine *h* (Biotium) in L-15 medium, without phenol red (Invitrogen), 10%FCS in the dark for 2 h before recording luminescence on the plate reader. After 10 seconds, cells were flashed with 470nm light (10^15^ photons) before immediately resuming recording for 60 seconds.

## Acknowledgements

We thank Andrew Gillis, Ariane Dimitris, Kasia Hammar, Elaine Seaver, Danielle de Jong, Paul Linser, Anne Zakrzewski, Michael Akam and Matthew Berriman for technical help and support. The research was supported by an EDEN NSF grant, Isaac Newton Trust grant, and by a Wellcome Trust Janet Thornton Fellowship (WT206194) to KR, a Natural Sciences and Engineering Research grant (A5056) to BKH and by a Biotechnology and Biological Sciences Research Council grant (BB/H006966/1) (F.L.) and a Leverhulme Trust grant (F/07 134/DA) to MT. MT is supported by European Research Council (ERC-2012-AdG 322790).

## Supplementary figures

**Supplementary figure 1:**
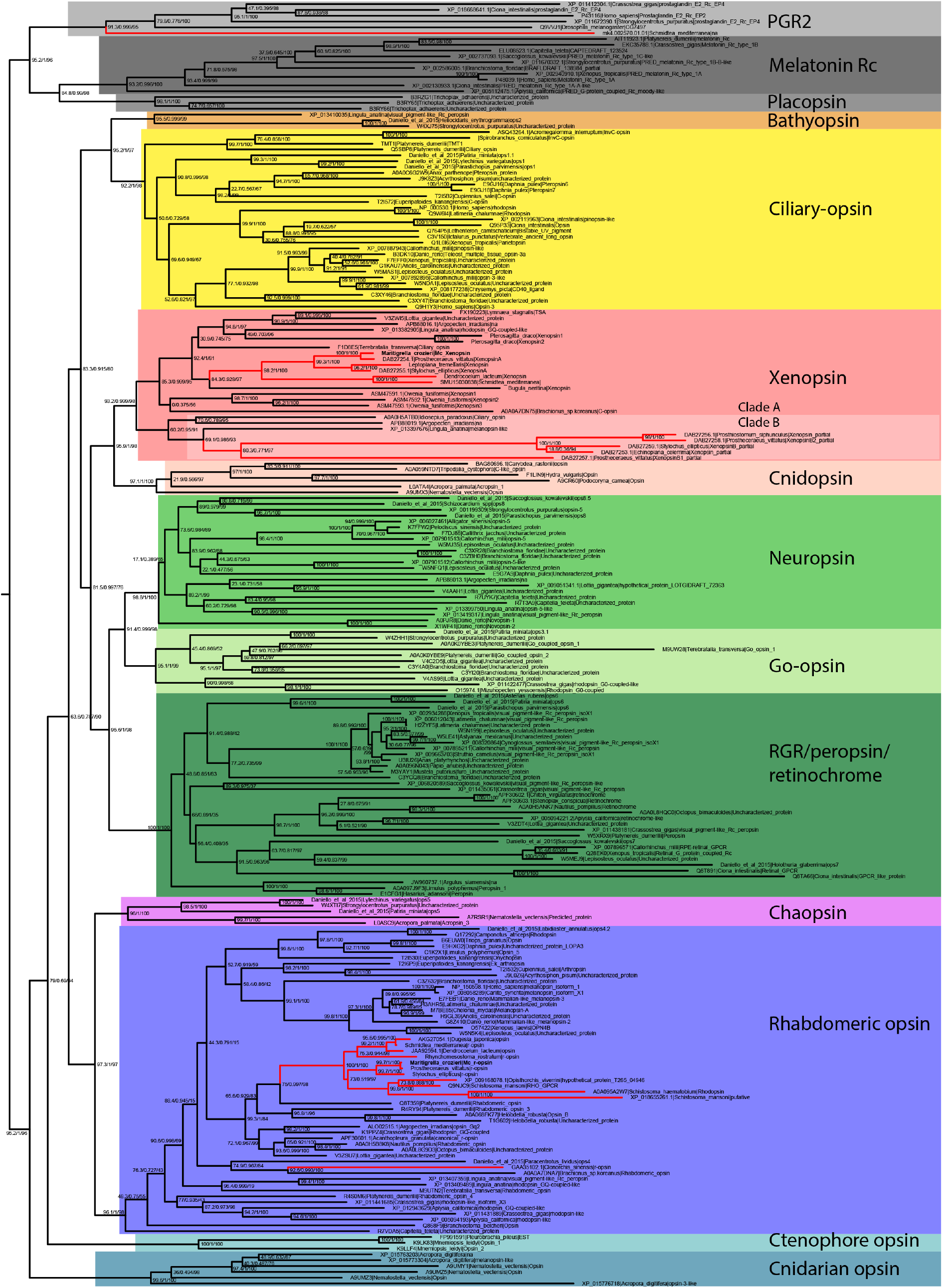
Uncollapsed tree of IQ-TREE phylogenetic reconstruction of opsin relationships. Node support values correspond to 1000 Ultrafast bootstrap replications, 1000 SH-aLRT replicates and approximate aBayes single Branch testing. Scale bar unit for branch length is the number of substitutions per site. Note low support for all deeper nodes including those uniting xenopsin/cnidopsin with tetraopsins, and the sister relationship between this clade and the c-opsins/bathyopsins. Branches in red correspond to flatworm opsin sequences.

**Supplementary figure 2:**
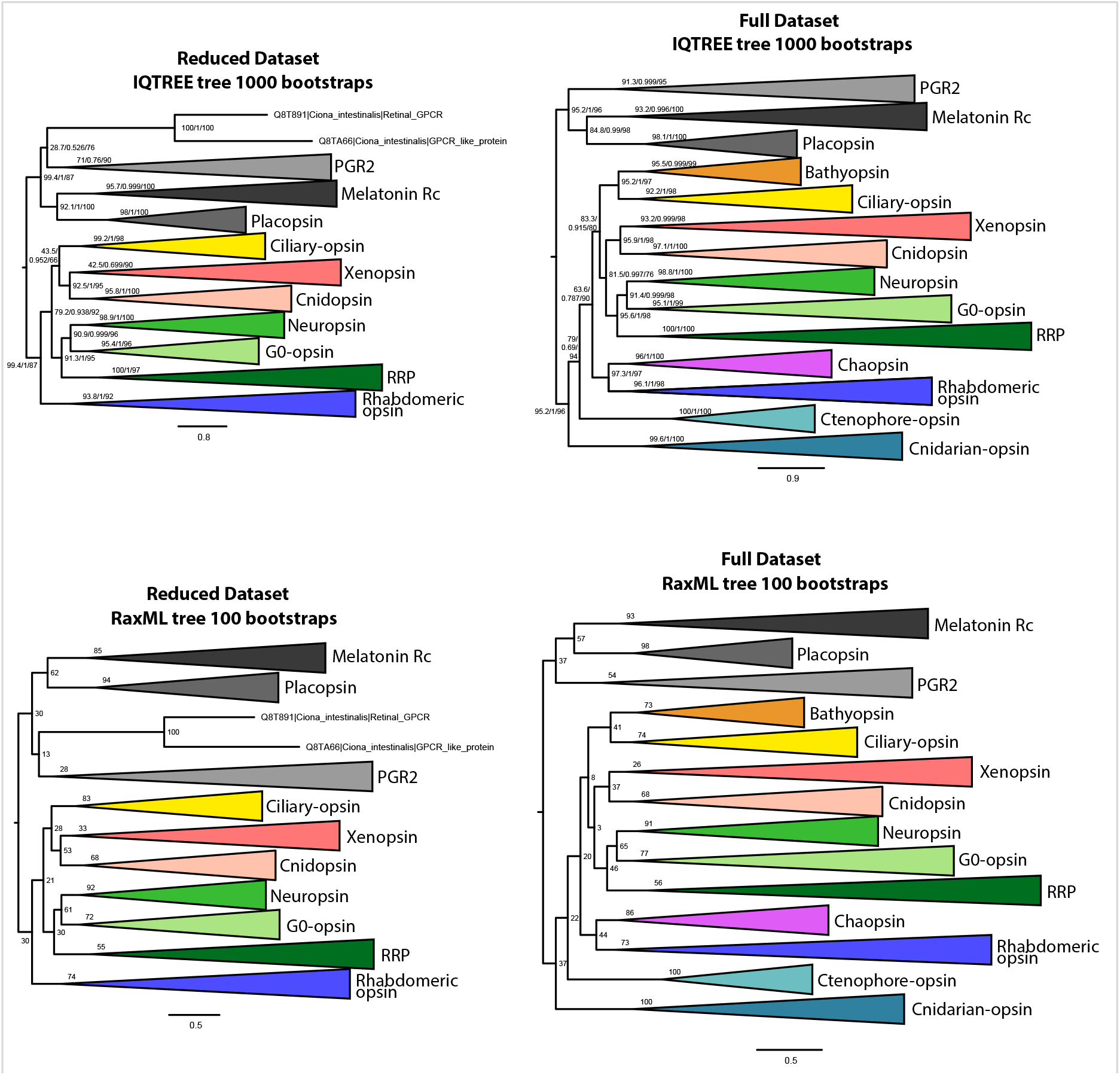
IQtree and RaxML trees showing the influence of the small opsin clades (i.e. chaopsins, bathyopsins, ctenophore and anthozoan opsins) on the position of xenopsins in relation to c-opsins and tetra-opsins (Neuropsin, Go-opsin and RRP); inclusion of these small opsin clades brings xenopsins close to tetraopsins (full dataset), their exclusion brings xenopsins close to c-opsins (reduced dataset).

**Supplementary figure 3:**
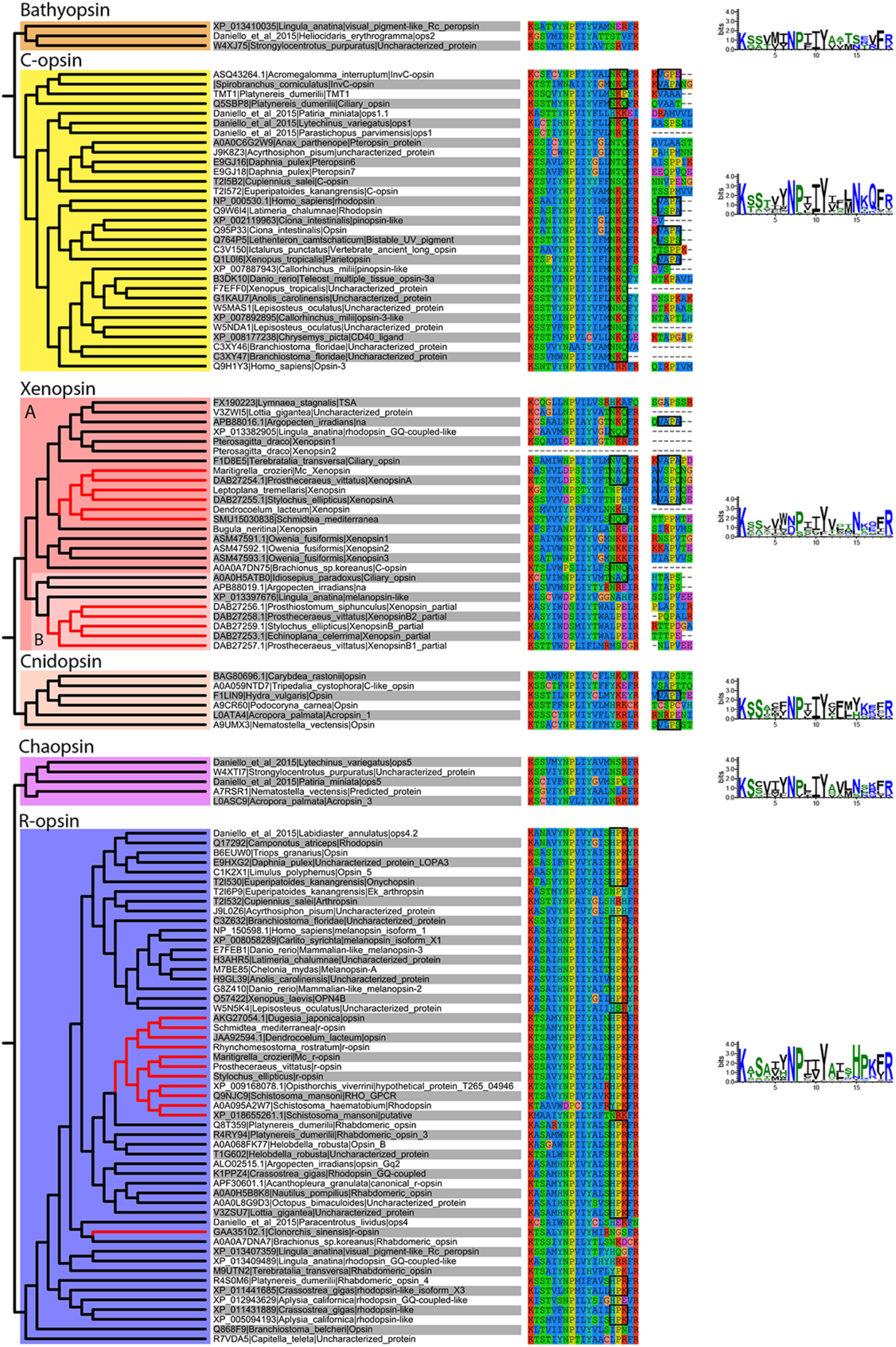
Alignment of major opsin clades showing conserved lysine in transmembrane domain VII, which binds to the retinal chromophore to form a photopigment. Some xenopsins possess a tripeptide motif, NxQ, which is also found in ciliary opsins and known to be crucial for G-protein activation. A number of flatworm xenopsin sequences in clade A have similar NxQ patterns (including *Maritigrella -* NAQ), while the motif differs considerably in the polyclad xenopsins of clade B, cnidopsins, bathyopsins and tetraopsins. An alignment of the C-terminal regions of ciliary opsins, xenopsins, cnidopsins, tetraopsins and bathyopsins shows, at a conserved position, similar VxPx motifs in flatworm clade A xenopsins (including *Mc xenopsin* - VSPQ) as well as a mollusk (*A. irradiens*) and brachiopod (*T. transversa*) xenopsin, it is also present in ciliary opsins from non-vertebrate chordates (tunicate and lamprey) and annelids, as well as in cnidopsin sequences. This motif binds the small GTPase Arf4 to direct vertebrate rhodopsin (a ciliary opsin) to the primary cilia. The presence of this motif in some ciliary opsins, xenopsins, cnidopsins may suggest a shared mechanism for the active delivery of these opsins to the cilia in CPRs.

**Supplementary figure 4:**
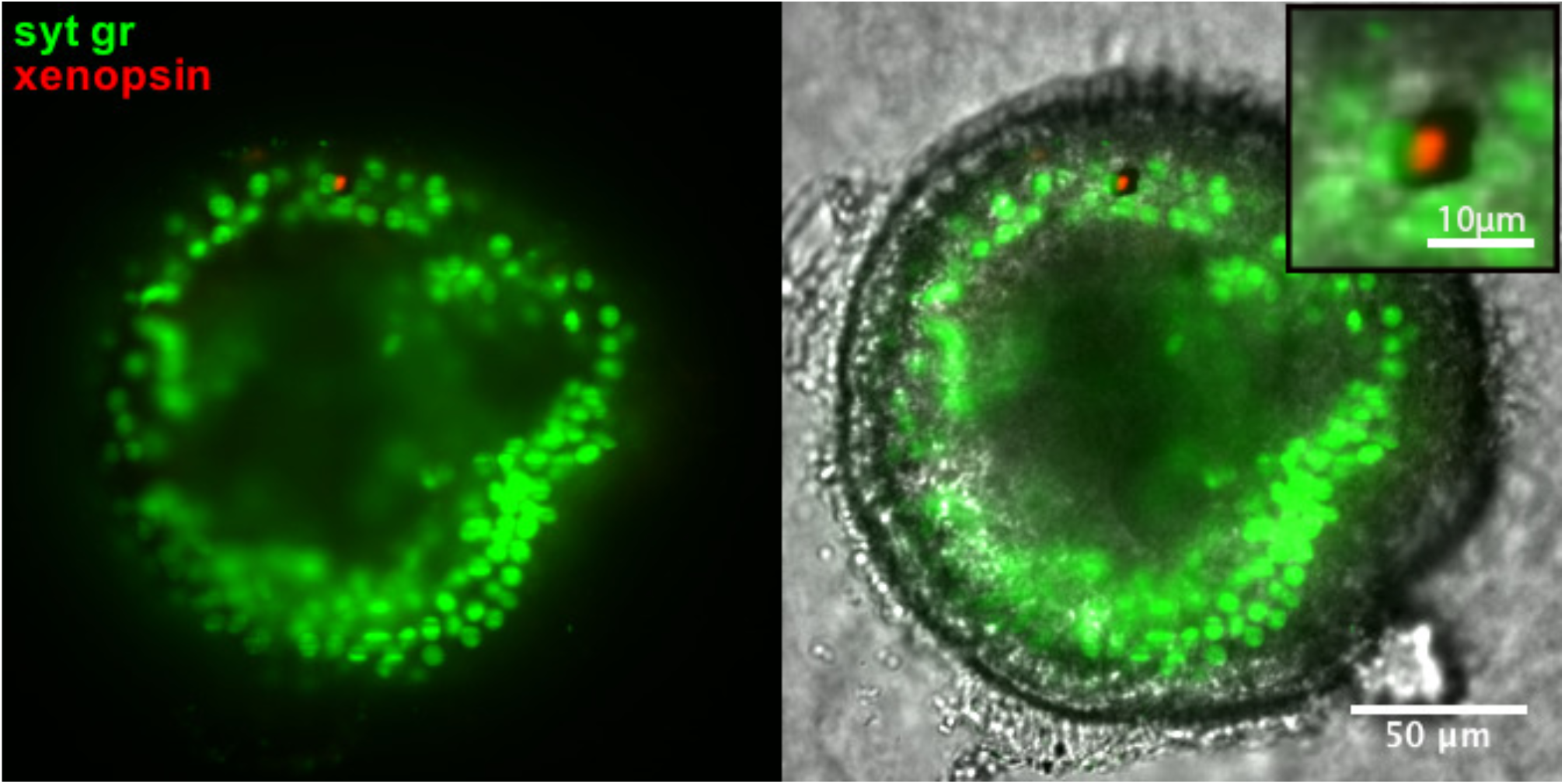
Xenopsin expression (red) in the epidermal eye during *Maritigrella crozieri* embryogenesis. The epidermal eye develops soon after gastrulation is complete and before development of the cerebral eyes. Syt gr = Sytox Green, staining nuclei. Bright-field also shows the photoreceptor pigments. Inset is a 3-time magnification.

**Supplementary figure 5:**
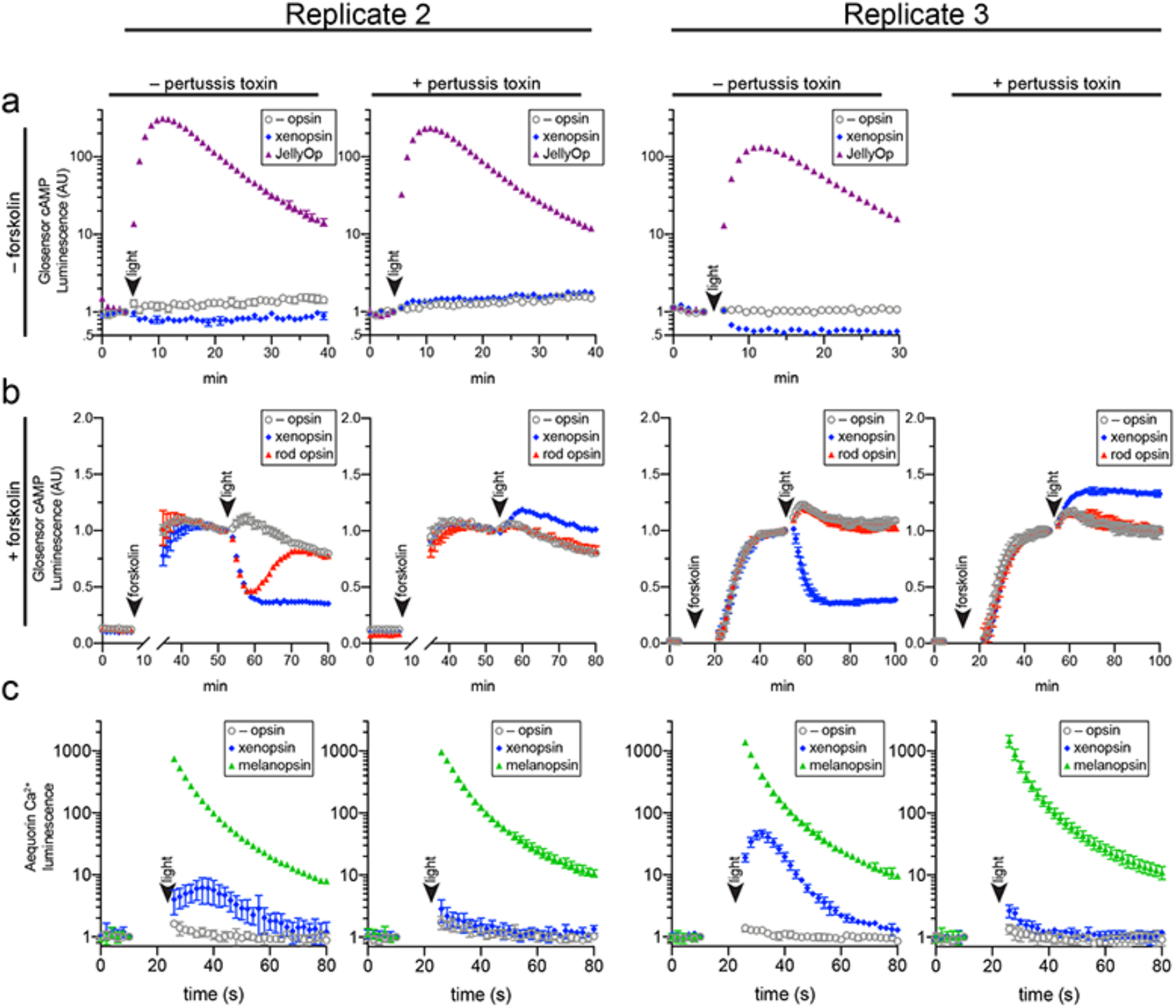
Two further biological replicates of the secondary messenger assays show there was substantial quantitative variation from day to day in the magnitude of responses to light and forskolin, but the qualitative response of each opsin was consistent (excluding one replicate in which rod opsin showed no activity, possibly due to a faulty preparation of plasmid DNA).

## Supplementary videos

**Supplementary video 1–.**
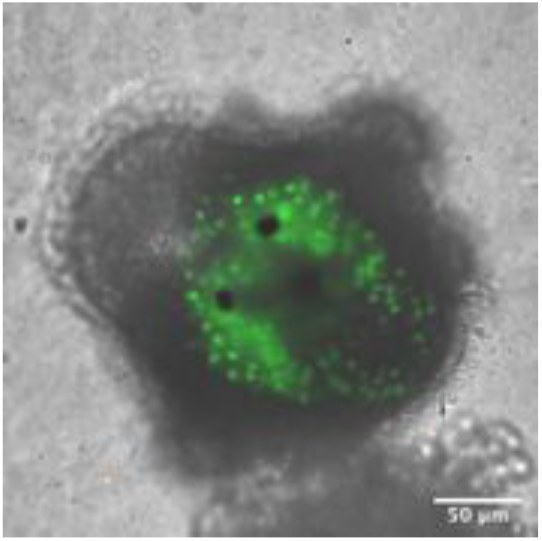
OpenSPIM z-stack through the anterior end of a *Maritigrella crozieri* larva showing xenopsin expression in the epidermal eye only. (Syt gr – Systox green nuclear stain, xenopsin – red).

**Supplementary video 2–.**
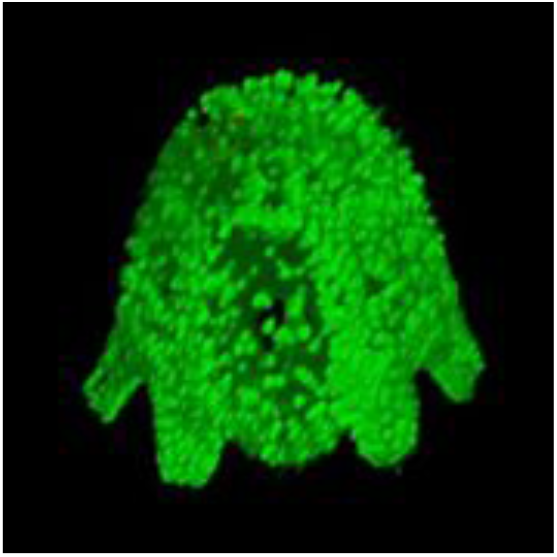
3D rendering of a multi-view OpenSPIM stack showing the dorso-lateral position of the xenopsin^+^ epidermal eye of the larval stage of *Maritigrella crozieri*. (Syt gr – Sytox green nuclear stain, xenopsin – red).

## Competing interests

On behalf of all authors I declare that there are no competing interests.

## References

The octopus genome and the evolution of cephalopod neural and morphological novelties. CB Albertin, O Simakov, T Mitros, ZY Wang, JR Pungor, E Edsinger-Gonzales, S Brenner, C Ragsdale, D Rokhsar (2015) Nature 524 (7564): 220–224. https://doi.org/10.1038/nature14668

Ciliary photoreceptors with a vertebrate-type opsin in an invertebrate brain. D Arendt, K Tessmar-Raible, H Snyman, AW Dorresteijn, J Wittbrodt (2004) Science 306:869–871. https://doi.org/10.1126/science.1099955

The enigmatic xenopsins. D Arendt (2017) eLife 6:e31781. https://doi.org/10.7554/eLife.31781

Reproducible and sustained regulation of Gαs signalling using a metazoan opsin as an optogenetic tool. Bailes HJ, Zhuang L-Y, Lucas RJ (2012) PLoS ONE 7(1): e30774. https://doi.org/10.1371/journal.pone.0030774

Human melanopsin forms a pigment maximally sensitive to blue light (λmax ≈ 479 nm) supporting activation of Gq/11 and Gi/o signalling cascades. H Bailes, R Lucas (2013). Proc R Soc B 280: 20122987.https://doi.org/10.1098/rspb.2012.2987

A live cell assay of GPCR coupling allows identification of optogenetic tools for controlling Go and Gi signaling. E Ballister, J Rodgers, F Martial, R Lucas (2018) BMC Biology 16:10 https://doi.org/10.1186/s12915-017-0475-2

Phototransduction in fan worm radiolar eyes. M Bok, M Porter, D Nilsson (2017) Current Biology 27 (14) R698–R699. https://doi.org/10.1016/j.cub.2017.05.093

Fine structure of the eyes of *Onithochiton neglectus* (Mollusca: Polyplacophora). PR Boyle (1956). Z Zellforsch 102:313–332.

TrakEM2 software for neural circuit reconstruction. A Cardona, S Saalfeld, J Schindelin, I Arganda-Carreras, S Preibisch, M Longair, P Tomancak, V Hartenstein, RJ Douglas (2012). PLoS ONE 7, e38011.

trimAl: a tool for automated alignment trimming in large-scale phylogenetic analyses. S Capella-Gutiérrez, JM Silla-Martínez, T Gabaldón (2009) Bioinformatics 25(15):1972–3.

Photoreceptors and photoreceptions in rotifers. P Clement, E Wurdak (1984). Photoreception and Vision in Invertebrates. 10.1007/978-1-4613-2743-1_8

The evolution of invertebrate photopigments and photoreceptors. TW Cronin, ML Porter (2014). Evolution of visual and non-visual pigments. (Marshall, Collins, eds). Springer USA.

Extraocular, non-visual, and simple photoreceptors: an introduction to the symposium. TW Cronin, S Johnsen (2016) Integr Comp Biol 56(5):758–763

Rhodopsin C terminus, the site of mutations causing retinal disease, regulates trafficking by binding to ADP-ribosylation factor 4 (ARF4). D Deretic, AH Williams, N Ransom, V Morel, PA Hargrave, A Arendt (2005) Proc Natl Acad Sci USA 102(9):3301. https://doi.org/10.1073/pnas.0500095102

Fine structure of the eyes of *Pseudoceros canadensis* (Turbellaria, Polycladida). RM Eakin, JL Brandenburger (1981) Zoomorphology 98:1–16.

A transcriptomic-phylogenomic analysis of the evolutionary relationships of flatworms. B Egger, F Lapraz, B Tomiczek, S Müller, C Dessimoz, J Girstmair, N Škunca, K Rawlinson, C Cameron, E Beli, M Antonio Todaro, M Gammoudi, C Noreña, MJ Telford (2015) Current Biology 25 (10) P1347–1353. https://doi.org/10.1016/j.cub.2015.03.034

Photoreceptors and photosensitivity in Platyhelminthes. Fournier A (1984). In: Ali M.A. (eds) Photoreception and Vision in Invertebrates. NATO ASI Series (Series A: Life Sciences), vol 74. Springer, Boston, MA DOI https://doi.org/10.1007/978-1-4613-2743-1_7

Light-sheet microscopy for everyone? Experience of building an OpenSPIM to study flatworm development. Johannes Girstmair, Anne Zakrzewski, François Lapraz, Mette Handberg-Thorsager, Pavel Tomancak, Peter Gabriel Pitrone, Fraser Simpson and Maximilian J. Telford. (2016) BMC Developmental Biology 16:22.10.1186/s12861-016-0122-0

Spectral tuning of phototaxis by a Go-opsin in the rhabdomeric eyes of platynereis. M Gühmann, H Jia, N Randel, C Verasztó, LA Bezares-Calderón, NK Michiels, S Yokoyama, G Jékely (2015) Current Biology 25:2265 2271. https://doi.org/10.1016/j.cub.2015.07.017

Melanopsin-containing retinal ganglion cells: architecture, projections, and intrinsic photosensitivity. S Hattar, HW Liao, M Takao, DM Berson, KW Yau (2002) Science 295(5557):1065-70. DOI:10.1126/science.1069609

Immunohistochemical (cLSM) and ultrastructural analysis of the central nervous system and sense organs in *Aeolosoma hemprichi* (Annelida, Aeolosomatidae). R Hessling, G Purschke (2000) Zoomorphology 120:65–78.

Presumed photoreceptive cilia in a ctenophore. GA Horridge (1964) Quart J Micr Sci 105(3): 311–317.

Cloning and sequence analysis of cDNA for the luminescent protein aequorin. S Inouye, M Noguchi, Y Sakaki, Y Takagi, T Miyata, S Iwanaga, FI Tsuji (1985) Proc. Natl Acad. Sci. USA 82, 3154–3158. 10.1073/pnas.82.10.3154

MAFFT multiple sequence alignment software version 7: improvements in performance and usability. K Katoh, DM Standley (2013) Mol Biol Evol 30(4):772–80.

Activation of transducin by bistable pigment parapinopsin in the pineal organ of lower vertebrates. Kawano-Yamashita E, Koyanagi M, Wada S, Tsukamoto H, Nagata T, Terakita A (2015) PLoS ONE 10(10): e0141280. https://doi.org/10.1371/journal.pone.0141280

Environmental stimuli, sense organs and behaviour in juvenile adult monogeneans. GC Kearn (1993) Bulletin Français de la Peche et de la Pisciculture 328:105–114.

Homologs of vertebrate Opn3 potentially serve as a light sensor in nonphotoreceptive tissue. M Koyanagi, E Takada, T Nagata, H Tsukamoto, A Terakita (2013) Proc Natl Acad Sci U S A 110(13):4998-5003. doi:10.1073/pnas.1219416110.

The brain and central nervous system of Müller’s larva. TC Lacalli (1983) Can J Zool 61:39–51.

Evolution of phototransduction, vertebrate photoreceptors and retina. T Lamb (2013) Progress in Retinal and Eye Research 36:52e119. 10.1016/j.preteyeres.2013.06.001

The ultrastructure of the eyes in larval and adult polyclads (Turbellaria). Alberto Lanfranchi, Celina Bedini and Enrico Ferrero. Hydrobiologia 84, 267–275 (1981). 0018-8158/81/0843-0267/$01.40.

Transcriptome analysis of the planarian eye identifies *ovo* as a specific regulator of eye regeneration. Lapan SW, Reddien PW (2012) Cell Reports 2:294–307. 10.1016/j.celrep.2012.06.018

Put a tiger in your tank: the polyclad flatworm *Maritigrella crozieri* as a proposed model for evo-devo. Lapraz F, Rawlinson KA, Girstmair J, Tomiczek B, Berger J, Jékely G, Telford MJ, Egger B. (2013) BMC Evo Devo 4:29. 10.1186/2041-9139-4-29

Nuclear genomic signals of the ‘microturbellarian’ roots of platyhelminth evolutinary innovation. Laumer CE, Hejnol A, Giribet G. (2015) eLife. 12;4. 10.7554/eLife.05503

Cd-hit: a fast program for clustering and comparing large sets of protein or nucleotide sequences. W Li, A Godzip (2006) Bioinformatics 22 (13) 1658–1659. 10.1093/bioinformatics/btl158

Sense organs of monogeneans. KM Lyons (1972) Zool. J. Linnean Soc. 51:181–199.

The BLAST sequence analysis tool. Madden T.L. (2002). In McEntyre, J. (ed.), The NCBI Handbook [Internet]. National Library of Medicine (US), National Center for Biotechnology Information, Bethesda, MD.

The amino terminus of the fourth cytoplasmic loop of rhodopsin modulates rhodopsin-transducin interaction. EP Marin, A Gopala Krishna, TA Zvyaga, J Isele, F Siebert, TP Sakmar (2000). The Journal of Biological Chemistry. 275, 1930–1936. 10.1074/jbc.275.3.1930

Photoreceptor number and outer segment disk membrane surface area in the retina of the rat: stereological data for whole organ and average photoreceptor cell. TM Mayhew, D Astle (1997). Journal of Neurocytology 26:53–61. https://doi.org/10.1023/A:1018563409196

Miller, M.A., Pfeiffer, W., and Schwartz, T. (2010) “Creating the CIPRES Science Gateway for inference of large phylogenetic trees” in Proceedings of the Gateway Computing Environments Workshop (GCE), 14 Nov. 2010, New Orleans, LA pp 1–8.

Single-photon sensitivity of lamprey rods with cone-like outer segments. A Morshedian, GL Fain (2015). Curr Biol 25, 484–487. 10.1016/j.cub.2014.12.031

Light adaptation and the evolution of vertebrate photoreceptors. A Morshedian, GL Fain (2017). J Physiol 595.14: 4947–4960. 10.1113/JP274211

Eye evolution: a question of genetic promiscuity. DE Nilsson (2004) Curr. Opin. Neurobiol 14, 407–414. 10.1016/j.conb.2004.07.004

The evolution of eyes and visually guided behavior. DE Nilsson (2009) Phil Trans R Soc B. 364:2833–2847.

Eye evolution and its functional basis. DE Nilsson (2013) Visual Neuroscience 30:5–20. https://doi.org/10.1017/S0952523813000035

A molecular analysis of neurogenic placode and cranial sensory ganglion development in the shark, Scyliorhinus canicular. O’Neill, RB McCole, CVH Baker (2007) Developmental biology 304 (1), 156-181 10.1016/j.ydbio.2006.12.029

Ciliary photoreceptors in the cerebral eyes of a protostome larva. YJ Passamaneck, N Furchheim, A Hejnol, MQ Martindale, C Lüter (2011) EvoDevo 2:6. https://doi.org/10.1186/2041-9139-2-6

Key transitions during the evolution of animal phototransduction: novelty, “tree-thinking,” co-option, and co-duplication David C. Plachetzki1 and Todd H. Oakley. Integrative and Comparative Biology, 47(5) 759–769. 10.1093/icb/icm050

Shedding new light on opsin evolution. ML Porter, JR Blasic, MJ. Bok, EG. Cameron, T Pringle, TW. Cronin, PR. Robinson (2012) Proc. R. Soc. B. 10.1098/rspb.2011.1819.

Ultrastructure of the phaosomous photoreceptors in *Stylaria lacustris* (Naididae, Oligochaeta, Clitellata) and their importance for the position of the Clitellata in the phylogenetic system of the Annelida. GJ Purschke (2003) Zool Syst Evol Res. 41:100–108. 10.1046/j.1439-0469.2003.00203.x

Photoreceptor cells and eyes in Annelida. G Purschke, D Arendt, H Hausen, MCM Muller (2006) Arthr Struct Dev. 35:211–230.

A gonad-expressed opsin mediates light-induced spawning in the jellyfish Clytia. G Quiroga Artigas, P Lapébie, L Leclère, N Takeda, R Deguchi, G Jékely, T Momose, E Houliston (2018) eLife 7:e29555 DOI:10.7554/eLife.29555

The last common ancestor of most bilaterian animals possessed at least 9 opsins. MD Ramirez, AN Pairett, MS Pankey, JM Serb, DI Speiser, AJ Swafford, TH Oakley (2016) Genome Biology and Evolution DOI:10.1093/gbe/evw248

Embryonic and post-embryonic development of the polyclad flatworm *Maritigrella crozieri*; implications for the evolution of spiralian life history traits. KA Rawlinson (2010) Front Zool.7:12. https://doi.org/10.1186/1742-9994-7-12

Ultrastructure of pigmented photoreceptors of larval *Multicotyle purvisi* (Trematoda, Aspidogastrea). K Rohde, NA Watson (1991) Parasitol Res. 77:485–490.

Software for bead-based registration of selective plane illumination microscopy data. Preibisch S, Saalfeld S, Schindelin J, Tomancak P. (2010) Nat Methods. 7:418–9. doi:10.1038/nmeth0610-418.

Efficient Bayesian-based multi-view deconvolution. Preibisch S, Amat F, Stamataki E, Sarov M, Singer RH, Myers E, Tomancak P (2014) Nat Methods. 11:645–8. doi:10.1038/nmeth.2929.

Double-stranded RNA specifically disrupts gene expression during planarian regeneration. A Sanchez-Alvarado, PA Newmark (1999) Proc Natl Acad Sci USA. 96:5049–5054. 10.1073/pnas.96.9.5049

Expression of ‘segmentation’ genes during larval and juvenile development in the polychaetes *Capitella* sp. I and *H. elegans*. EC Seaver, LM Kaneshige (2006) Dev Biol. 289:179-194. 10.1016/j.ydbio.2005.10.025

Comparative morphology of photoreceptors in free-living plathelminths - a survey. B Sopott-Ehlers (1991) Hydrobiologia 227:23l–9.

Photoreceptors in species of the Macrostomida (Plathelminthes): ultrastructural findings and phylogenetic implications. B Sopott-Ehlers, W Salvenmoser, D Reiter, R Rieger, U Ehlers. (2001) Zoomorphology. 121:1–12.

Switch of rhodopsin expression in terminally differentiated *Drosophila* sensory neurons Sprecher, Simon G and Desplan, Claude (2008) Nature 454. 533 https://doi.org/10.1038/nature07062

RAxML version 8: a tool for phylogenetic analysis and post-analysis of large phylogenies. A Stamatakis (2014). Bioinformatics 30(9):1312-3. doi:10.1093/bioinformatics/btu033.

Disc morphogenesis in vertebrate photoreceptors. RH Steinberg, SK Fisher, DH Anderson (1980) J. Comp. Neurol. 190:501–518. http://dx.doi.org/10.1002/cne.901900307

The opsins. A Terakita (2005) Genome Biol. 6:213. 10.1186/gb-2005-6-3-213

Stacked endoplasmic reticulum sheets are connected by helicoidal membrane motifs. M Terasaki, T Shemesh, N Kasthuri, RW Klemm, R Schalek, KJ Hayworth, AR. Hand, M Yankova, G Huber, JW Lichtman, TA Rapoport, MM Kozlov (2013). Cell 154, 285–296. 10.1016/j.cell.2013.06.031

Melatonin signaling controls circadian swimming behavior in marine zooplankton. MA Tosches, D Bucher, P Vopalensky, D Arendt (2014) Cell 159:46–57. https://doi.org/10.1016/j.cell.2014.07.042

W-IQ-TREE: a fast online phylogenetic tool for maximum likelihood analysis. J Trifinopoulos, LT Nguyen, A von Haeseler, BQ Minh. (2016) Nucleic Acids Research 44, W232– W235, https://doi.org/10.1093/nar/gkw256

Diversity and functional properties of bistable pigments. Hisao Tsukamoto and Akihisa Terakita (2010) Photochem. Photobiol. Sci. 9:1435–1443 DOI:10.1039/c0pp00168f

A ciliary opsin in the brain of a marine annelid zooplankton is ultraviolet-sensitive, and the sensitivity is tuned by a single amino acid residue. H Tsukamoto, IS Chen, Y Kubo, Y Furutani (2017) Journal of Biological Chemistry 292:12971–12980. 10.1074/jbc.M117.793539

Co-expression of xenopsin and rhabdomeric opsin in photoreceptors bearing microvilli and cilia. O Vöcking, I Kourtesis, SC Tumu, H Hausen (2017). eLife 6:e23435. https://doi.org/10.7554/eLife.23435

Ciliary and rhabdomeric photoreceptor-cell circuits form a spectral depth gauge in marine zooplankton. C Verasztó, M Gühmann, H Jia, VB Veedin Rajan, LA Bezares-Calderón, C Piñeiro-Lopez, N Randel, R Shahidi, NK Michiels, S Yokoyama, K Tessmar-Raible, G Jékely (2018) eLife 7:e36440 DOI:10.7554/eLife.36440

Seasonal changes in the lower jaw skeleton in male Atlantic salmon (*Salmo salar* L.): remodelling and regression of the kype after spawning. PE Witten, BK Hall (2003) J Anatomy.203: 435–450.

Fine structure of a potential photoreceptor organ in the larva of *Bugula neritina* (Bryozoa). RM Woollacott, RL Zimmer (1972) Z. Zellforsch. 123,458–469. doi:10.1007/BF00335542

Phototransduction Motifs and Variations. KW Yau, RC Hardie (2009). Cell 139 246–264

Molecular evidence for convergence and parallelism in evolution of complex brains of cephalopod molluscs: insights from visual systems. M. A. Yoshida, A. Ogura, K. Ikeo, S. Shigeno, T. Moritaki, G. C. Winters, A. B. Kohn and L. L. Moroz (2015). Integrative and Comparative Biology 55(6)1070–1083. doi:10.1093/icb/icv049

The repertoire of G protein-coupled receptors in the human parasite *Schistosoma mansoni* and the model organism *Schmidtea mediterranea*. M Zamanian, MJ Kimber, P McVeigh, SA Carlson, AG Maule, TA Day (2011). BMC Genomics 12:596. 10.1186/1471-2164-12-596

